# Adaptive Remodeling of the MPXV B21R Receptor-Binding Domain Enhances DC-SIGN Interaction and Identifies Conserved CTL Targets for T-Cell Vaccine Development

**DOI:** 10.64898/2026.03.02.708970

**Authors:** Swatantra Kumar, Arpita S. Harnam, Saurabh Kumar, Janusz T. Paweska, Ahmed S. Abdel-Moneim, Shailendra K. Saxena

## Abstract

Global Mpox transmission is imposing public health concern as the number of cases is progressively increasing since its first major outbreak in 1996. Therefore, understanding its global epidemiological transformation and its underlying mechanism is crucial to decipher the immune evasion strategies exhibited by recent MPXV strains. In the present study, we analyzed the trend of global Mpox epidemiology and identified the current multinational outbreak which has initiated in 2017 from Africa. To explore the molecular basis of this transformation, we considered the B21R protein of MPXV as it may have played a role in viral adaptation and immune escape mechanism as one of the important MPXV structural proteins. Our data shows that Mpox has significantly transformed from 1996 to 2025, where MPXV strains from 2022, 2023, and 2024 are closely clustered whereas 2025 is closely related to 2017 MPXV strain. Structural modeling of B21R using AlphaFold uncovers a modular architecture comprising a putative receptor-binding N-terminal region (p-RBD), a central ectodomain, and a membrane-anchored C-terminal segment. Mapping solvent accessibility across the full-length B21R protein revealed that p-RBD exhibited highest solvent exposure compared to other B21R protein domains. As a potential cellular receptor for entry into the host targeted cell, we evaluated the interaction of p-RBD of B21R protein with CRD region of DC-SIGN, which showed the gradual increase in the binding affinity with acquired mutations. Moreover, we found alteration in the O-linked glycosylation sites at p-RBD regions of B21R protein which is crucial for the MPXV entry into the host cell. Importantly, we observed significant changes in linear B cell epitopes of p-RBD, impacting the humoral immunity, while CTL epitopes remained conserved. Hence, we showed the significance of B21R p-RBD as a T-cell based vaccine candidate for prevention of Mpox. This study provides novel insights into the recent global transmission of the Mpox and explored a plausible mechanism of humoral immune escape strategies through progressive mutations in the B21R protein and potential development of T-cell based vaccine candidate.

**Significance statement:** Our study represents the global transmission dynamics of Mpox and immune evasion strategy of recent MPXV strains. Epidemiological transformation analysis revealed that the current multinational outbreak of Mpox originated in Africa in 2017, highlighting the expanding global footprint of MPXV. Our analysis based on the B21R protein shows the evolutionary adaptation of the MPXV associated with progressive mutations responsible for increased affinity towards the DC-SIGN receptor and potential reason for increased infectivity. Importantly, alterations in O-linked glycosylation sites and linear B cell epitopes show potential antigenic drift in the recent Mpox outbreaks showing immune escape strategies. These findings provide insights into Mpox epidemiology and the molecular basis of MPXV adaptation, informing vaccine design, therapeutic strategies, and improved countermeasures against future outbreaks.

**Graphical Abstract:** 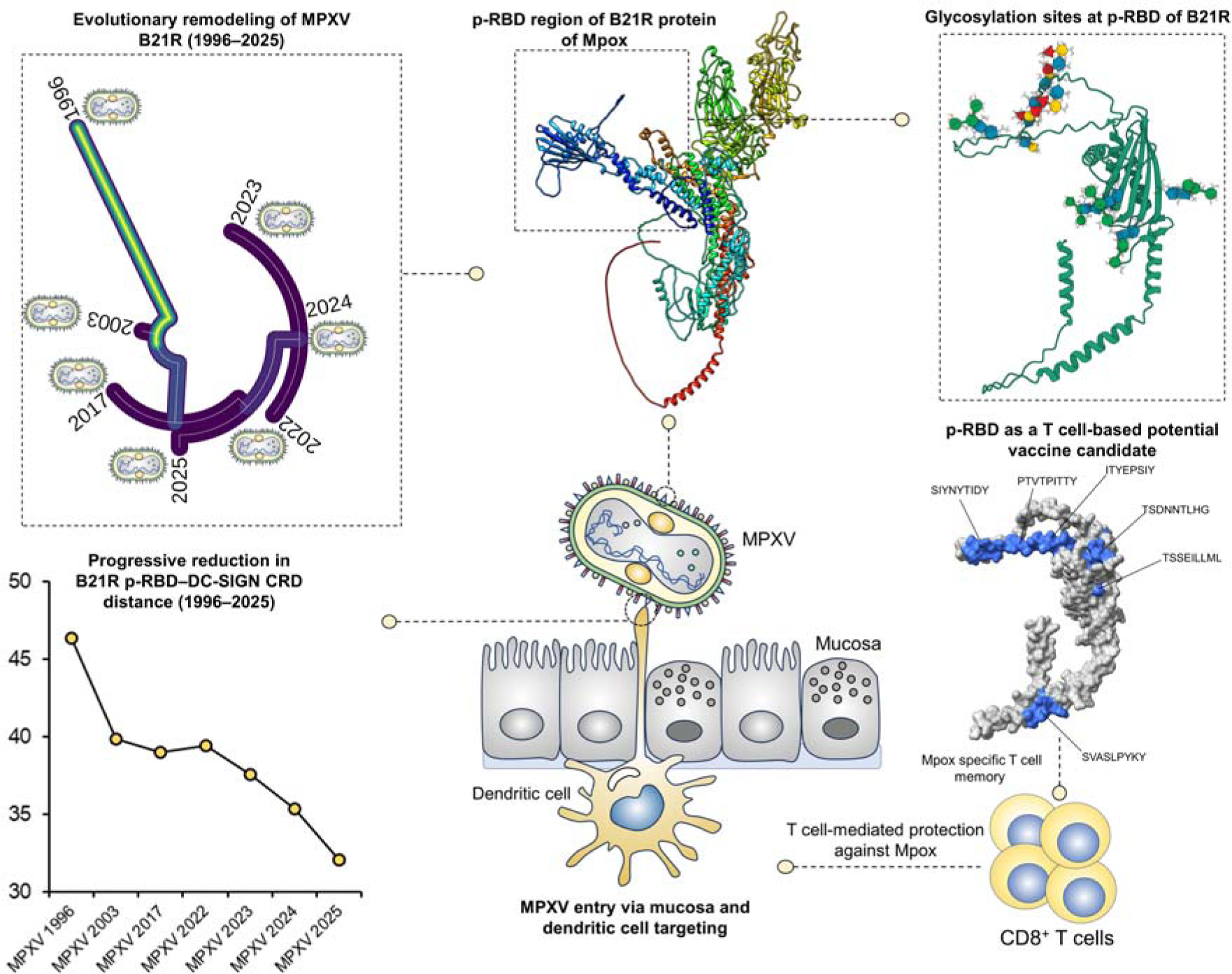

## Introduction

Mpox is an infectious disease, earlier known as monkeypox, caused by the monkeypox virus (MPXV) which belongs to a genus *Orthopoxvirus* and *Poxviridae* family [1, 2]. MPXV infected individuals may experience fever, headache, myalgia, fatigue, painful rashes/mucosal lesions and enlarged lymph nodes. Mpox is mostly a self-limiting infection wherein the symptoms may last for 2-3 weeks. Most of the infected individuals fully recovered and the case fatality rate is around 3%-6% [3]. Transmission of MPXV is through close contact with someone infected with MPXV [4]. The first sporadic cases of Mpox were reported in the 1970s-1990s in Western Africa and Central Africa. The first major outbreak of Mpox was reported in Zaire in 1996 with 71 human cases [5]. Subsequently, Mpox resurgence was reported in 2000-2004 in Democratic Republic of Congo. Notably, the first outbreak of Mpox outside Africa was reported in the United States in 2003 [6]. Importantly, a large outbreak was further reported in Nigeria in 2017 [7]. Recently, due to a global multinational outbreak of Mpox reported from year 2022, imposing immense challenges due to significant increase in the number of cases worldwide. [8].

Phylogenetically there are two characterized clades of MPXV clade I and clade II which have been further divided into two subclades [9]. Importantly, people infected with clade I were at the higher risk of mortality due to more severe infection as compared with the people infected with clade II [10]. Prominently, the Clade IIb strains of MPXV were responsible for the recent global outbreak of Mpox which started in the year 2022 [11]. However, recent cases of Mpox in Democratic Republic of the Congo (DRC) are due to clade I strains and therefore caused ∼ 47,000 Mpox cases and more than ∼1,000 deaths. The clade I MPXV strains have been transmitted to some of the neighboring countries including Central African Republic, Burundi, Rwanda and Uganda. However, travel-associated cases of Mpox were reported in Kenya, Zambia, Zimbabwe, Germany, United Kingdom, Thailand and India [12]. On August 14, 2024, the WHO declared Mpox as the public health emergency of international concern [13]. As the current transmission dynamics of Mpox impose a global public health concern, it is imperative to understand the current global epidemiological transformation in recent Mpox outbreaks and its impact on the global population in terms of transmissibility and immune evasion strategies of emerging MPXV variants.

MPXV can bind to host cell surface molecules like GAGs, chondroitin sulfate or integrin for internalization [14]. MPXV can enter the host cell either directly by fusion with the host plasma membrane or by endocytosis. Once inside the cell, the MPXV coat is removed and its genome is released in cytoplasm [15]. MPXV gene expression is tightly regulated, early genes are expressed soon after virus entry. These early genes code for proteins involved in immune evasion and replication. Intermediate genes also code for proteins involved in virus replication. Late expressing genes encode for proteins involved in viral assembly [16, 17]. The virus is assembled as immature virions (IMVs), which later on transform into mature virions (MVs) which may be released by cell lysis. MV undergoes further wrapping in Golgi and transforms into enveloped virions (EEVs). EEV can leave the infected cell either by forming the actin tail assembly or by budding [16] **(Figure 1)**.

**Figure 1.**
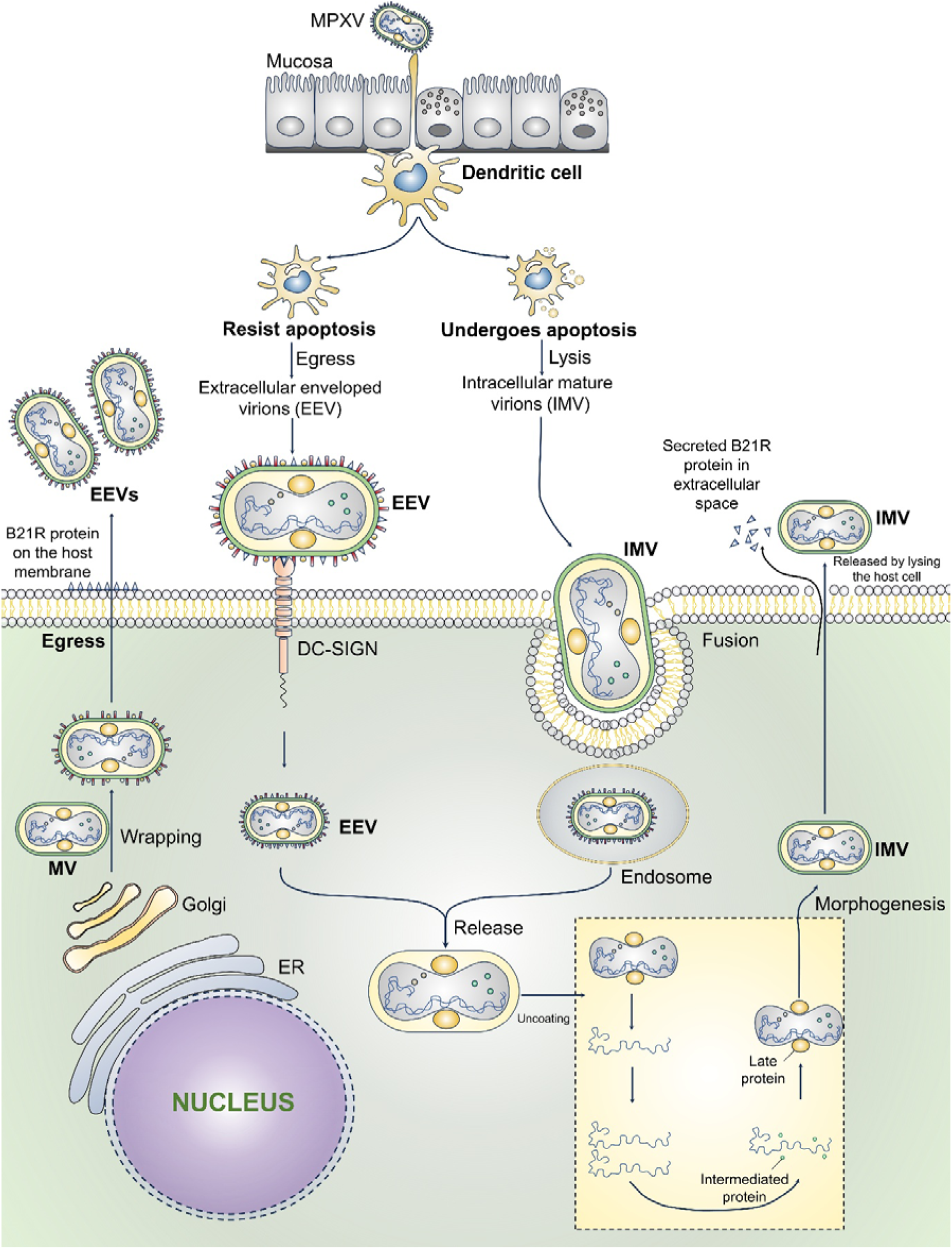
Proposed role of B21R during MPXV infection and dissemination. Schematic representation the role of B21R glycoprotein in viral entry, immune interaction, and dissemination. B21R is most likely involved in the entry of the MPXV via interacting with DC-SIGN receptor. Dendritic cell interacts with MPXV infection when infection initiate with mucosal epithelium. MPXV infection can lead to apoptosis leading to intracellular mature virions (IMVs) release. However, resistance of apoptosis results into the release of extra-cellular enveloped virions (EEVs). B21R on the viral envelope and host cell membranes is suggested to mediate viral attachment to DC-SIGN receptors on dendritic cells of EEVs. Whereas, IMVs enter host cells through membrane fusion or endocytosis. Both IMV and EEV can undergo uncoating accompanied by cytoplasmic replication, including early, intermediate, and late gene expression. Viral assembly results in the generation of mature virions (MVs) which can be membrane wrapped by the Golgi-ER network to generate EEVs via taking the B21R on their envelope during egress process. In addition, B21R protein secreted into the extracellular space along with the IMVs if the target cell is undergoing apoptosis. This conceptual model highlights the functional importance of B21R and its putative receptor-binding domain (p-RBD) as a target for immunological intervention and T-cell based vaccine design.

In the present study, we analyzed the transformation of global epidemiology of Mpox during its major outbreaks from the year 1996 to 2025 and analyzed mutational mapping of B21R protein in major and current Mpox outbreaks considering its importance in emergence of B.1 lineage of Mpox for the current outbreaks [18]. B21R is a surface glycoprotein which has been identified as one of the important antibody targets with various immunodominant epitopes [19, 20]. B21R is present only in MPXV thus it could help to distinguish MPXV from other Orthopoxviruses [5]. It can suppress T-cell activation by altering TCR signaling thus lowering the expression of IL-2 and INF-γ and impair the function of both CD4^+^ and CD8^+^ cells [21]. Mutations in B21R can alter the host interaction and viral transmission [22]. As of now the B21R structure has not been characterized. Therefore, we determined B21R structure and analyzed its putative receptor binding domain (p-RBD) along with its affinity towards carbohydrate recognition domain (CRD) of a potential cell surface receptor Dendritic Cell-Specific Intercellular adhesion molecule-3-Grabbing Non-integrin (DC-SIGN) also known as CD209 [23]. DC-SIGN is a C-type lectin receptor on immature dendritic cells (DC) in peripheral tissues, it can interact with glycosylated viral proteins and can facilitate the viral entry and dissemination. Not always as a direct entry receptor, DC-SIGN enables the virus to undergo trans-infection, where the virus is captured by DCs and subsequently transferred to permissive cells enhancing the virus spread to secondary lymphoid organs [24]. Importantly, it has been shown that MPXV viral messenger RNA present in DC-SIGN^+^ cells in the upper dermis, epithelial cells [25]. Therefore, assessing the affinity of B21R towards the DC-SIGN may reveal the viral transmission and immune modulation. Moreover, we analyzed the N and O-linked glycosylation sites and antigenic epitopes of B21R proteins from the major Mpox outbreak to understand its further role in global epidemiological transformation of Mpox. Understanding the B21R protein structure and function can help in developing effective vaccines as it is an important antibody target comprising key immune-dominant epitopes.

## Results

### Global epidemiology transformation showing multinational outbreak of Mpox

To understand the transformation of global epidemiology of Mpox, we have analyzed the continent wise trend of Mpox outbreaks based on the number of confirmed cases on a 7 days average basis relative to the population per one million cases. Continent wise cases of Mpox have been highlighted in the continent maps where a darker colour represents the higher number of cases in that region **(Figure 2A)**. Global trends in the current Mpox outbreak peaked in the year 2022 in the western regions including North America, South America and Europe. Whereas, the number of cases in this region declined significantly in 2023,2024 and 2025. Importantly, the Mpox cases sporadically peaked in the year 2023 in Asia and later in 2025. However, the cases in Asia were significantly less in the year 2022 and 2024 in comparison with 2023 and 2025. Upon analyzing the trend of Mpox cases in Africa, we have found four periodic year wise peaks identified in the year 2022, 2023, 2024 and 2025 **(Figure 2B)**. However, the significant rise in the Mpox cases in the Africa region started in 2024 and peaked in 2025. Moreover, the Mpox cases in Oceania peaked in the year 2022 and in 2024 with some cases in 2025. Importantly, when we tracked the Mpox cases from the year 2003, we have found that Mpox originated from Africa and subsequently transmitted to Europe and Asia and later resulted in the multinational outbreaks of Mpox **(Figure 2C)**.

**Figure 2.**
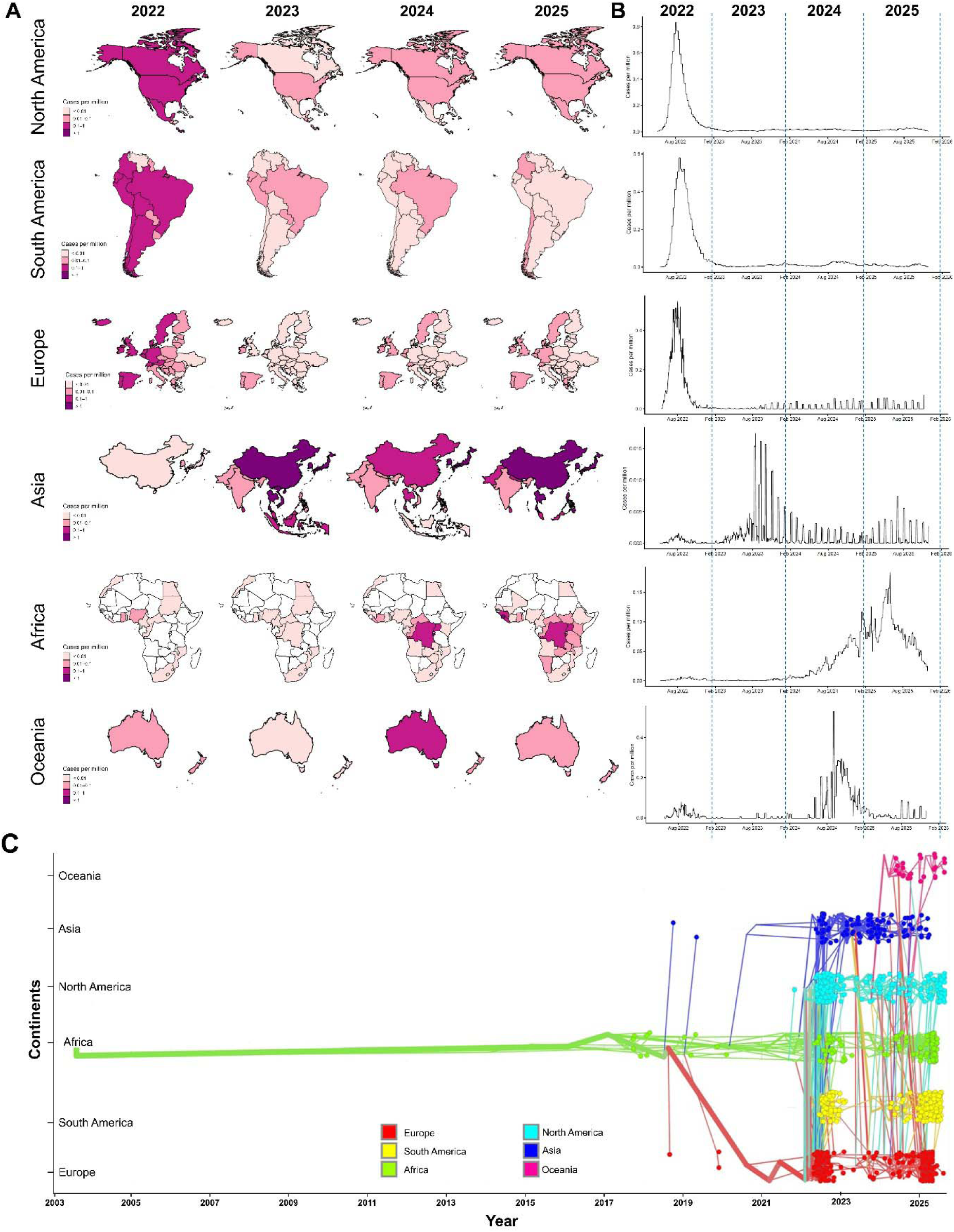
Global spatiotemporal dynamics of Mpox incidence across major continents during recent multinational outbreak. **A**. Continent wise incidence of Mpox (cases per million population) from 2022 to 2025. Choropleth maps showing the burden of mpox in each region of North America, South America, Europe, Asia, Africa, and Oceania. Color intensity corresponds to the intensity of incidence where darker colors showing high case rates. The maps highlight the widespread outbreak during 2022 followed by heterogeneous regional persistence and resurgence patterns till 2025. **B.** Temporal trends in mpox incidence (cases per million population) across continents between 2022 and 2025. The line plots showing continent-specific epidemic patterns with an initial peak of the outbreak in 2022 and further decreasing incidence rates in most regions and in the later stages, local outbreaks surge in Asia, Africa, and Oceania. Vertical dashed lines indicate transitions between calendar years. **C.** Distribution of reported mpox cases between 2003 and 2025 on a long-term continental basis. Every point is an indicator of reported cases allocated to a continent and connecting lines show that some temporal transitions occurred in the geographic reporting. Tracking of the cases showing that the Mpox originated from Africa and subsequently transmitted to Europe and Asia and later resulted in the multinational outbreaks of Mpox from 2022-2025 outbreak.

### MPXV strains of multinational outbreaks are closely linked together in the phylogenetic analysis

To understand the phylogenetic relationship among the recent MPXV strains, we have retrieved the OPG210 protein (homolog of B21R) sequence of MPXV strains from different continents and analyzed the phylogenetic relationship among them **(Table S1)**. All the sequences were phylogenetically analyzed and the consensus sequences were used for the multiple sequence alignment (**Figure S1, S2**). We performed the continent wise phylogenetic analysis B21R protein sequences based on the consensus sequences from 2022, 2023, 2024 and 2025 **(Figure 3A)**. We have found the most divergent MPXV is in the year 1996 and the recent multinational outbreaks are closely clustered. The final transformed tree is unrooted and has 7 tips, with 12 total nodes. The tree is fully bifurcating except for the root node, which has 3 offspring which are closely clustered. The total length of the tree (sum of all branch lengths) is 0.01382; the tree is not clock-like: the minimum height of a leaf is 0.0006125, while the maximum height is 0.008715. Branch lengths of the tree range between 0 and 0.008715, with mean 0.001256 and median 0.000509 (89% highest-density interval: 0-0.008715, 95% HDI: 0-0.008715) **(Figure 3B)**.

**Figure 3.**
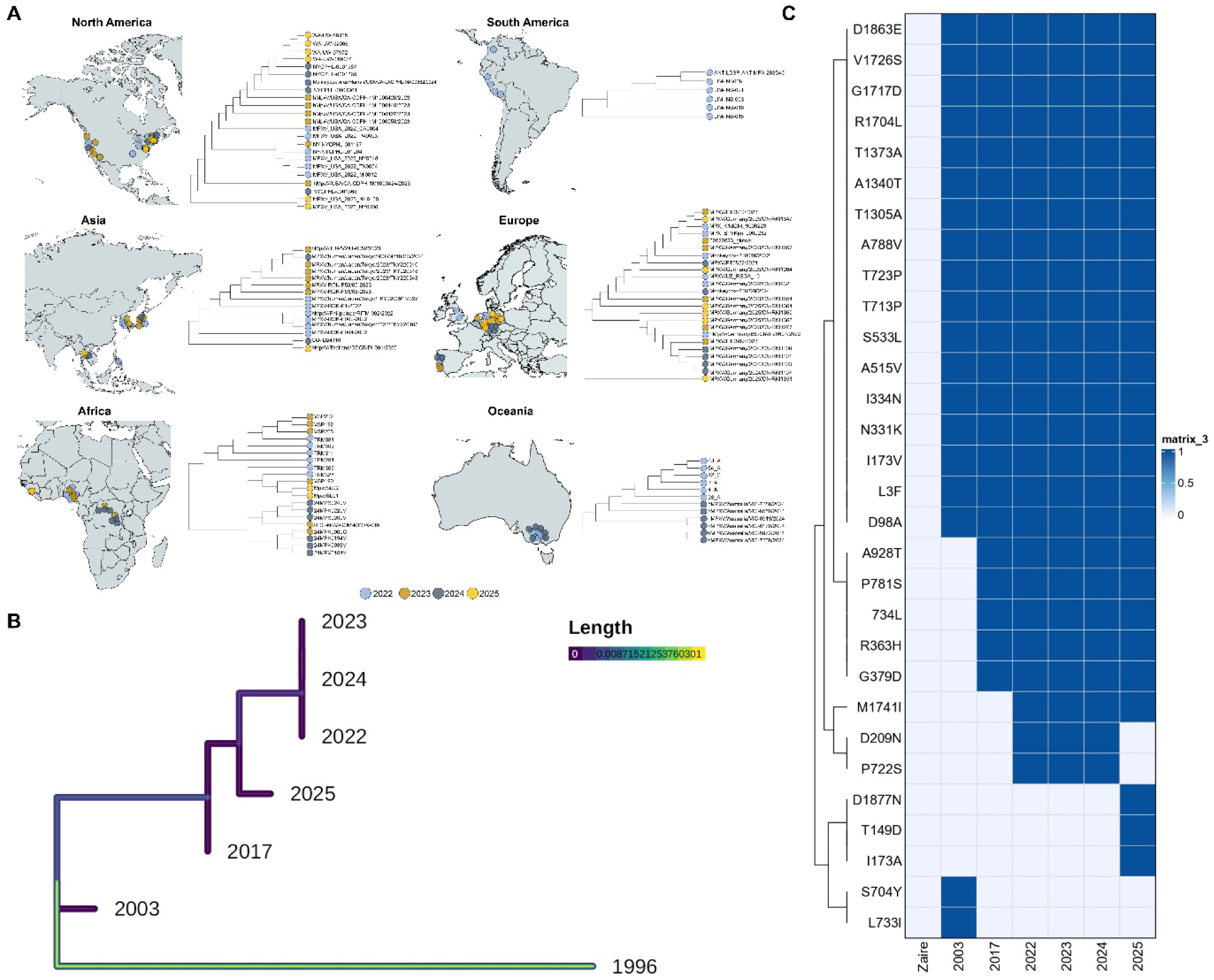
Phylogeographic distribution and mutational landscape of MPXV B21R protein sequences. **A.** Geographical maps of continents showing different regions collection of MPXV B21R sequences from 2022 to 2025. The continent wise phylogenetics showing the consistency of B21R protein sequence across the collection year which are represented as a unique color code. **B.** Based on the consensus year wise MPXV sequence, time-scaled phylogenetic tree from 1996 to 2025 showing the most divergent MPXV is in the year 1996 and the recent multinational outbreaks are closely clustered. The evolutionary divergence is equivalent to the length of branches. **C.** Heatmap showing the amino acid substitutions in the B21R protein in MPXV lineages from 1996 to 2025 where Zaire is representing the MPXV strain from 1996. Individual amino acid replacements are represented in rows and MPXV lineages are represented by columns. Darker shading indicates presence of mutations, whereas lighter shading indicates absence.

### Mutational hotspot in B21R protein in the recent MPXV strains

To understand the mutational hotspots of recent MPXV strains in B21R protein, we first compared the MPXV strains from the major outbreaks with the reference strain of MPXV Zaire-96-I-16. Importantly, we have identified 19 mutations in the 2003 strain as compared to the Zaire-96-I-16 strain. Prominently, 17 of these mutations as L3F, D98A, I173V, N331K, I334N, A515V, S533L, T713P, T723P, A788V, T1305A, A1340T, T1373A, R1704L, G1717D, V1726S, D1863E were found in rest of the strains of MPXV in subsequent major outbreaks. However, 2 mutations such as S704Y and L733I were unique to the 2003 strain of MPXV. Moreover, 2017 strains of MPXV exhibited 5 novel mutations as R363H, G379D, L734, P781S, A928T, which were subsequently found in the current Mpox outbreaks 2022, 2023, 2024, and 2025 strains. In addition to these previous mutations, three new mutations as M1741I, D209N, P722S, appeared in the 2022 MPXV strain and persisted in the 2023 and 2024 strains. However, the 2025 strain carries only the M1741I mutation, with the other two absent, and additionally acquired two new unique mutations as T149D and D1877N were identified. **(Figure 3C)**.

### Structure of B21R and its putative receptor binding region

To understand the role of B21R protein in the transmissibility of the recent Mpox outbreaks, we generated the structure of B21R protein through AlphaFold server. The model generated through the AlphaFold showed a pTM score of 0.53 where a score above 0.5 means the overall predicted fold for the complex might be like the true structure. Upon structural analysis we have found that B21R protein is a “Y” shaped tertiary structure composed of putative receptor binding regions and a membrane anchored transmembrane region **(Figure 4A)**. Based on the predicted aligned error plot (PAE) where the X-axis represents the Scored Residue and the Y-axis represents the Aligned Residue of the B21R protein and the diagonal line represents self-alignment. Moreover, we have found that B21R structure exhibits well defined regions with flexible loops enabling it to interact with the potential cellular receptor. The main diagonal of this plot is dark in color which suggests that the AlphaFold has higher confidence in intra-domain residue positioning. The lighter color represents sites which may be due to flexibility or lack of structural constraints **(Figure 4B)**.

**Figure 4.**
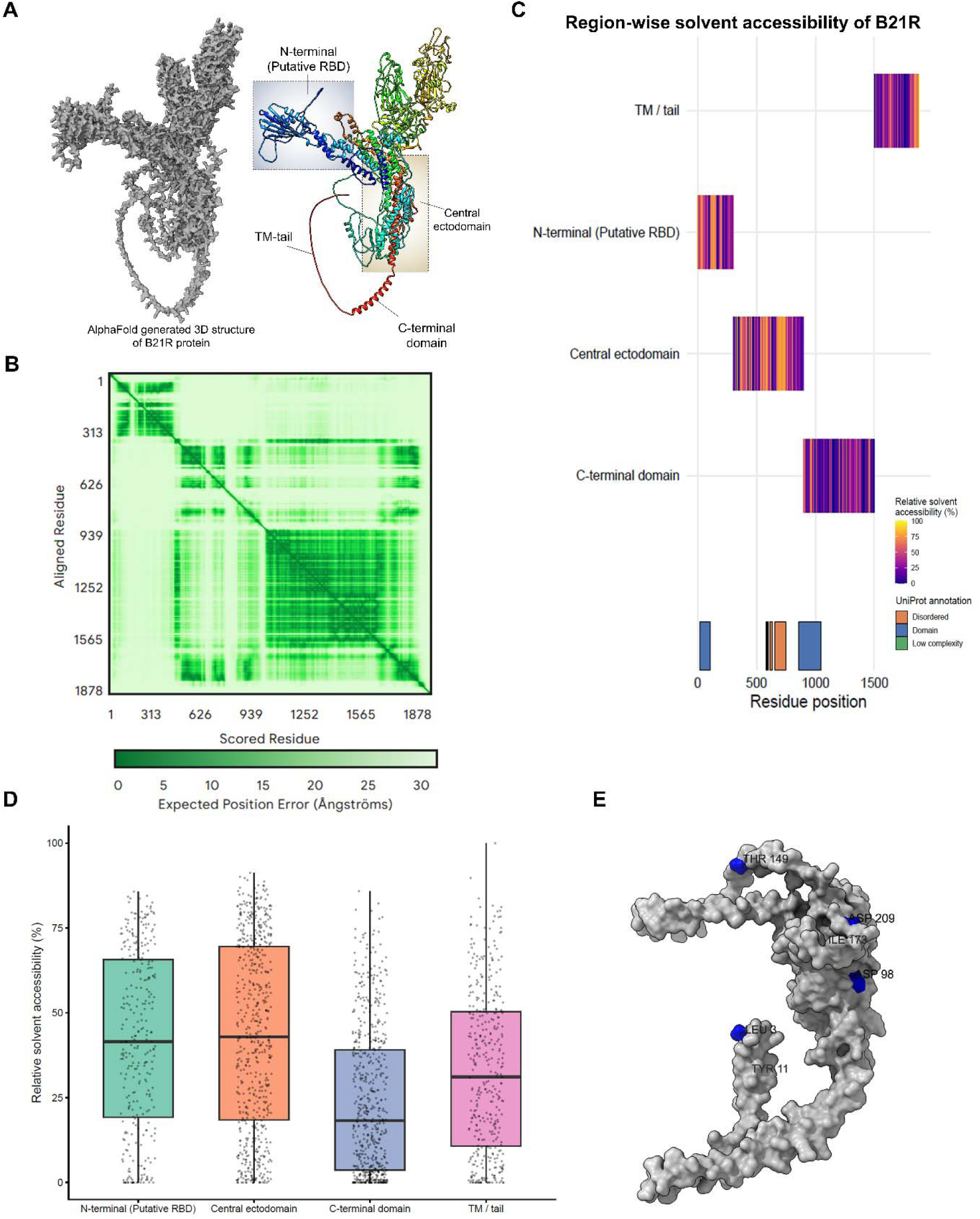
Structural organization and solvent accessibility analysis of the MPXV B21R protein. **A.** AlphaFold generated three-dimensional structure of MPXV B21R protein. The full-length model (left) and domain-annotated structure (right) are showing the N-terminal region (putative receptor-binding domain, p-RBD), central ectodomain, C-terminal domain and transmembrane (TM) tail. **B.** Predicted aligned error (PAE) plot showing the confidence in the B21R protein tertiary model where the X and Y axis represents the position of the residues and the color scale represents the predicted error distance. Lower values (darker green) indicate higher confidence in residue-residue positioning, whereas lighter regions represent increased structural uncertainty. The map showing overall structural consistency of the ectodomain regions including predicted p-RBD. **C.** Region-wide solvent accessibility profile of the B21R protein showing relative solvent accessibility (%) consistency predominantly in N-terminal p-RBD and central ectodomain regions which are relatively overlapping with the UniProt annotations. **D.** Box-and-scatter plots showing that the N-terminal p-RBD and central ectodomain have greater solvent exposure than the C-terminal domain and TM-tail region and indicating that these regions are more readily accessible by immune recognition. **E.** Surface representation of the B21R structure highlighting solvent-exposed residues within the p-RBD region.

### Region wise solvent accessibility of B21R revealed highly exposed N-terminal domain

SASA serve as a critical structural determinant that can reveal the rationale for the B21R p-RBD domain involved in evolution of Mpox. Mutations in the solvent exposed residues in viral surface glycoprotein are most likely to be responsible for the immune escape mechanisms. Therefore, we mapped relative solvent accessibility (RSA) of the residues over the entire B21R protein and compared the region-wise solvent accessibility across the annotated structural regions that revealed significant heterogeneity **(Figure 4C)**. As compared with the other region of the B21R protein, the p-RBD and central ectodomain region exhibited the higher solvent accessibility, showing the surface exposed architecture. Whereas, the transmembrane region and the tail regions showed variable accessibility due to partial membrane localization and regional conformational adaptability. The global Kruskal–Wallis test was performed to compare the significance of the solvent accessibility across the B21R protein regions. p-RBD region showed highest median solvent accessibility followed by the central ectodomain region **(Figure 4D)**. These findings support the structural observation that the p-RBD region is substantially surface exposed, contributing to its possible functional activity in receptor binding and immune recognition.

The solvent-accessible residues in the N-terminal domain were plotted onto the AlphaFold-predicted B21R structure to place solvent-exposed residues in the three-dimensional space. Even though several amino acid replacements were observed in the B21R sequence, a few of the mutations have been chosen to be examined further in terms of their structural implications. The repeated residues were observed over various outbreak years, were localized to the N-terminal portion, and had high solvent accessibility. Notably, overlay of the mutations on the residues L3, D98, T149, I173 and D209 on the AlphaFold-predicted structure showed that they were concentrated on a continuous patch of the surface, suggesting that they did not occur due to random genetic drift but they played a common functional role **(Figure 4E)**. On the other hand, mutations found in buried areas or other structurally constrained regions were less probable to influence interactions with hosts. Collectively, these observations indicate that mutations exposed to the surface are a functionally interesting hot spot, which is typical of a p-RBD.

### Higher affinity of recent MPXV B21R protein towards CRD domain of DC-SIGN

Structure of putative receptor binding domain was generated for all the major outbreaks from 1996 to 2025 for which the best clusters were selected based on the van der Waals energy and electrostatic energy **(Figure 5A)**. These structures were then subjected to their affinity and distance from the CRD domain of DC-SIGN. The CRD domain is located at the distal end of the extracellular portion of the DC-SIGN which adopts C-type lectin fold, comprising a core with conserved calcium-binding sites critical for glycan recognition exhibited by crucial residues as E347, N349, E354, N365 and D366 form coordinates with calcium ions for carbohydrate binding and provides structural support for glycan binding **(Figure 5B)**.

**Figure 5.**
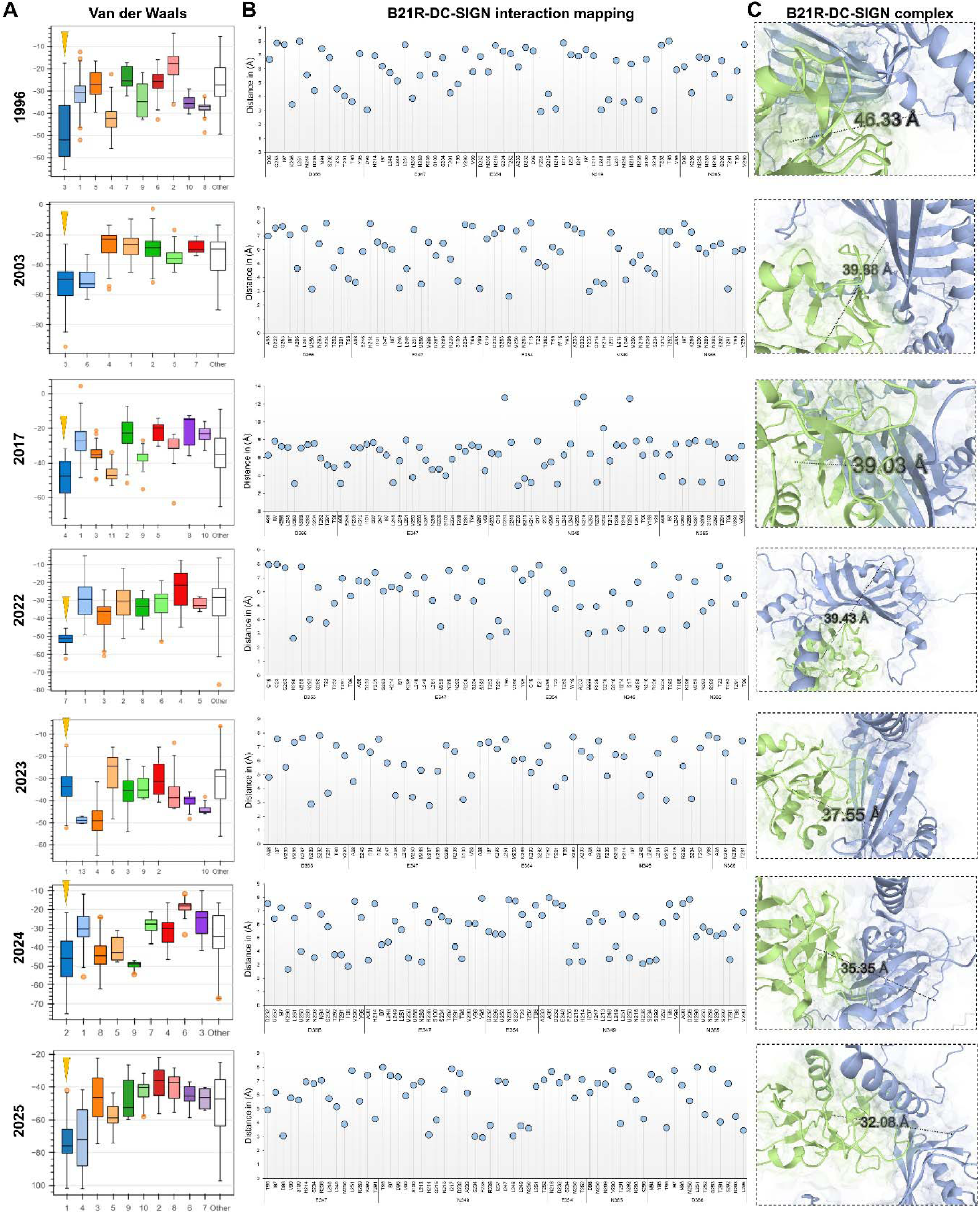
Structural interaction analysis of MPXV B21R with DC-SIGN across evolutionary lineages of Mpox. **A.** van der Waals interaction showing the clusters of the interaction and the distance between putative receptor of B21R protein with CRD region of DC-SIGN receptor across the MPXV strains from 1996 to 2025 where the most significant clusters identified as reverse yellow arrowheads. Overall interaction energies remained stable across time, suggesting conserved receptor-binding potential. **B.** Scatter plots showing intermolecular distances between interacting residues of representative 1996, 2003, 2017, 2022, 2023, 2024 and 2025 MPXV strains and the CRD region of DC-SIGN. The conserved interface regions are found in all lineages indicating that receptor-binding geometry is stable in the B21R p-RBD region. **C.** Molecular interaction of B21R (blue) and CRD region of DC-SIGN (green) suggesting progressive structural stabilization while maintaining conserved receptor engagement where intermolecular distances gradually decrease.

We have found that the distance between p-RBD and the CRD domain of DC-SIGN of Zaire-96-I-16 was 46.3Å which was found to further reduced in 2003 outbreak showed a distance of 39.88Å from CRD region, 2017 showed 39.03Å, 2022 showed 39.43Å, 2023 showed 37.55Å, 2024 showed 35.35Å and 2025 showed 32.08 Å **(Figure 5C)**. Upon analyzing the contact map of the putative receptors of B21R protein derived from the major outbreaks of Mpox with the CDR region of DC-SIGN receptor **(Table S2)**, we have found that the affinity of B21R is highest for 2025. Moreover, recent outbreak strains showed progressively more favorable binding free energies (ΔG) and reduced dissociation constants (KD). Importantly, the 2025 MPXV strain exhibited significant enhancement in the overall intermolecular contacts, which suggests that the electrostatic and hydrophobic interactions are optimized at the binding interface **(Table S3)**. This finding supports higher affinity of B21R protein with the CRD region of DC-SIGN may be responsible for the gradual increase of the global Mpox cases.

### Variation in the O-linked glycosylation at p-RBD of B21R protein in recent MPXV strains

B21R protein utilizes the heparan sulfate-like glycans for the attachment to the host cells; we further analyzed and mapped the N-and O-linked glycosylation in the B21R protein. These interactions rely on the sulfation patterns and structural features of the heparan sulfate glycans present on B21R protein. N-glycosylation involves the attachment of glycans to the nitrogen atom of asparagine (Asn) residues in proteins and involves immune escape strategy. O-glycosylation involves the attachment of glycans to the oxygen atom of serine (Ser) or threonine (Thr) residues in proteins and involves entry of the virus. Therefore, we performed the mapping of N-linked and O-linked glycans and analyzed the changes among the 1996, 2003, 2017, 2022, 2023, 2024 and 2025 **(Table S4, Table S5)**. The predicted N-linked and O-linked glycosylation sites were monitored to represent both conservation and variability, which allowed a comparison of the MPXV lineages with the help of heatmap representations. Interestingly, we have found that the N-linked glycosylation sites remain consistent across the MPXV strains. However, the O-linked glycosylation sites were transformed across the strains **(Figure 6A)**. To represent the glycosylation observed in the heatmap, we performed the mapping of the N and O-linked sites on the p-RBD **(Figure 6B)**. We have observed that the N-linked sites were widely distributed across the p-RBD and therefore involved in the protein stability and immune shielding. However, O-linked sites were found to be localized in the flexible loop region of p-RBD, representing its role in the host receptor engagement. Furthermore, the molecular representation of these sites was analyzed in detail **(Figure 6C)**. The proxy glycans were conjugated to the predicted N-linked and O-linked residues with representative units of monosaccharides, such as N-acetylglucosamine, mannose, galactose, and fucose. These models indicate that O-linked glycans are positioned to directly influence surface topology and accessibility of epitopes, and N-linked glycans are more stable and scaffold shapes. These findings taken together are consistent with a model of conserved N-linked glycosylation in immune evasion and strain-specific O-linked glycosylation in antigenic variation and different host interactions.

**Figure 6.**
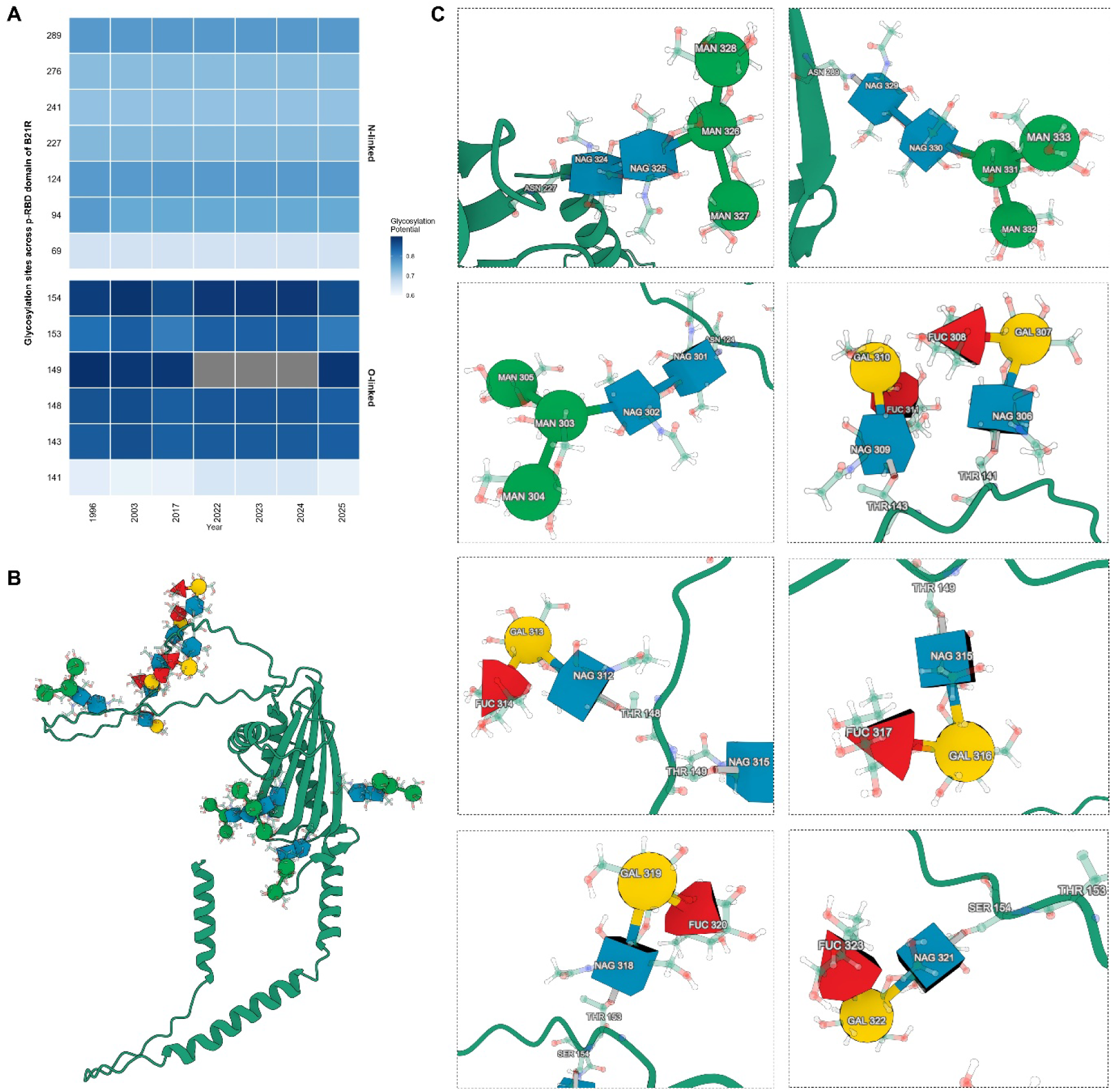
Glycosylation landscape of the MPXV B21R p-RBD region. **A.** Heatmap of the predicted N-linked and O-linked glycosylation sites across B21R p-RBD region of selected representative MPXV strains from 1996 to 2025. The intensity of color depicts the glycosylation potential scores. Some of the glycosylation sites are also preserved across lineages, while variations observed at selected O-linked positions, indicating largely stable glycosylation patterns during MPXV evolution. **B.** Structural mapping of predicted glycosylation sites onto the AlphaFold-predicted three-dimensional structure of the B21R protein. These glycans are located across exposed regions of the ectodomain, including the putative receptor-binding region, suggesting potential roles in immune recognition and protein stability. **C.** Detailed structural views of representative glycan moieties attached to B21R residues. N-acetylglucosamine (NAG), mannose (MAN), galactose (GAL), and fucose (FUC) residues are shown forming predicted glycan chains at specific glycosylation sites. The glycans are predominantly located on solvent-exposed surfaces, indicating accessibility to host immune components.

### Immune escape strategy due to antigenic variation in recent MPXV strains

To understand the immune escape strategies employed by the recent MPXV strains, we determined the antigenic variations among the MPXV strains via comparing linear B cell epitopes and CTL epitopes across the MPXV strains. We have found that there has been no significant change in the CTL epitopes among the 1996, 2003, 2017, 2022, 2023 2024 and 2025 MPXV strains **(Table S6)**. CD8+ T-cell epitopes showed high level of conservation among all the MPXV strains, with no substantial gain or loss of dominant CD8^+^ T-cell epitopes during evolution, showing the consistency in cellular immune response mediated by B21R protein. In contrast, we have found significant variation in the linear B cell epitopes specifically concentrated at the p-RBD region which is crucial for the interaction of the B21R protein with the DC-SIGN receptor **(Table S7)**. Heatmap-based comparison of epitope scores across outbreak years demonstrated dynamic remodeling of B-cell epitope landscapes, with several regions exhibiting fluctuating antigenicity scores and altered epitope boundaries in recent strains **(Figure 7A)**. Notably, epitope segments spanning residues 26–93, 104–164, and 194–211 showed consistent divergence in epitope strength and sequence composition between early (1996–2003) and recent (2022–2025) MPXV strains. Structural mapping of these variable B-cell epitopes on the three-dimensional B21R models indicated that antigenic alterations occur mostly in solvent-exposed surface loops and β-strands of the p-RBD region **(Figure 7B-H)**. Although the fundamental structure of the receptor-binding area is preserved, the reconstruction of surface-exposed B cell epitopes of recent MPXV strains suggests adaptive modulation of humoral immune recognition across the MPXV strains from 1996-2025.

**Figure 7.**
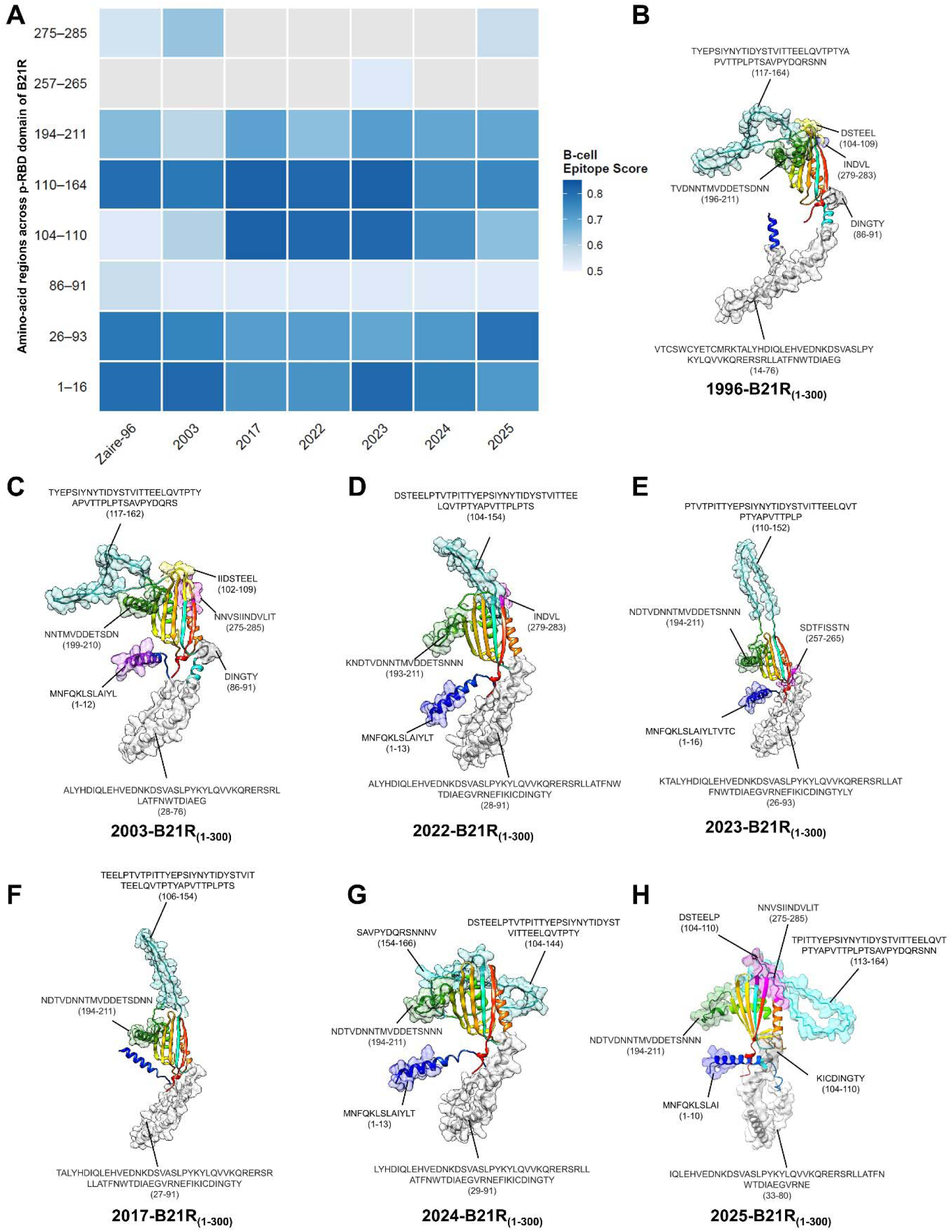
Predicted B-cell epitope landscape of the MPXV B21R p-RBD region. **A.** Heatmap showing predicted linear B-cell epitope scores across defined amino acid regions of the B21R p-RBD domain across MPXV strains from 1996 to 2025. Color intensity represents the B-cell epitope prediction score, indicating significant variation across the p-RBD regions. **B-H.** Structural mapping of predicted B-cell epitopes onto AlphaFold-derived models of the B21R p-RBD region for representative MPXV isolates from 1996 (B), 2003 (C), 2017 (F), 2022 (D), 2023 (E), 2024 (G), and 2025 (H). Predicted epitope regions are highlighted in distinct colors and mapped onto the protein surface. The epitopes cluster predominantly within the putative receptor-binding domain (p-RBD) and adjacent solvent-exposed loops. The spatial distribution of predicted epitopes shows substantial structural conservation across MPXV lineages, with minimal positional shifts despite sequence variation.

### B21R p-RBD harbors conserved CTL epitopes with MPXV T-cell vaccine potential

B21R p-RBD was analyzed using PRIME v2.1 to identify various 9-mer peptides with high predicted binding by MHC class I and favorable TCR recognition scores in representative HLA alleles (HLA-A01:01, HLA-A25:01, HLA-B07:02, and HLA-B18:01). A number of peptides were found %Rank_bestAllele below 0.1%, representing highest immunogenicity. The highest-ranking epitopes included ITTYEPSIY, PTVTPITTY, SIYNYTIDY, and TSEEILLML which localized in a continuous area of the p-RBD creating an immunogenic hot spot **(Figure 8A)**. The sequence alignment of MPXV strains (1996-2025) showed that these predicted CTL epitopes, including the key HLA anchor residues, are fully conserved, in spite of the occurrence of recurrent mutations in the neighboring solvent-exposed areas **(Figure 8B)**. This is indicative of structural or functional restriction maintaining T-cell epitope integrity in the p-RBD.

**Figure 8.**
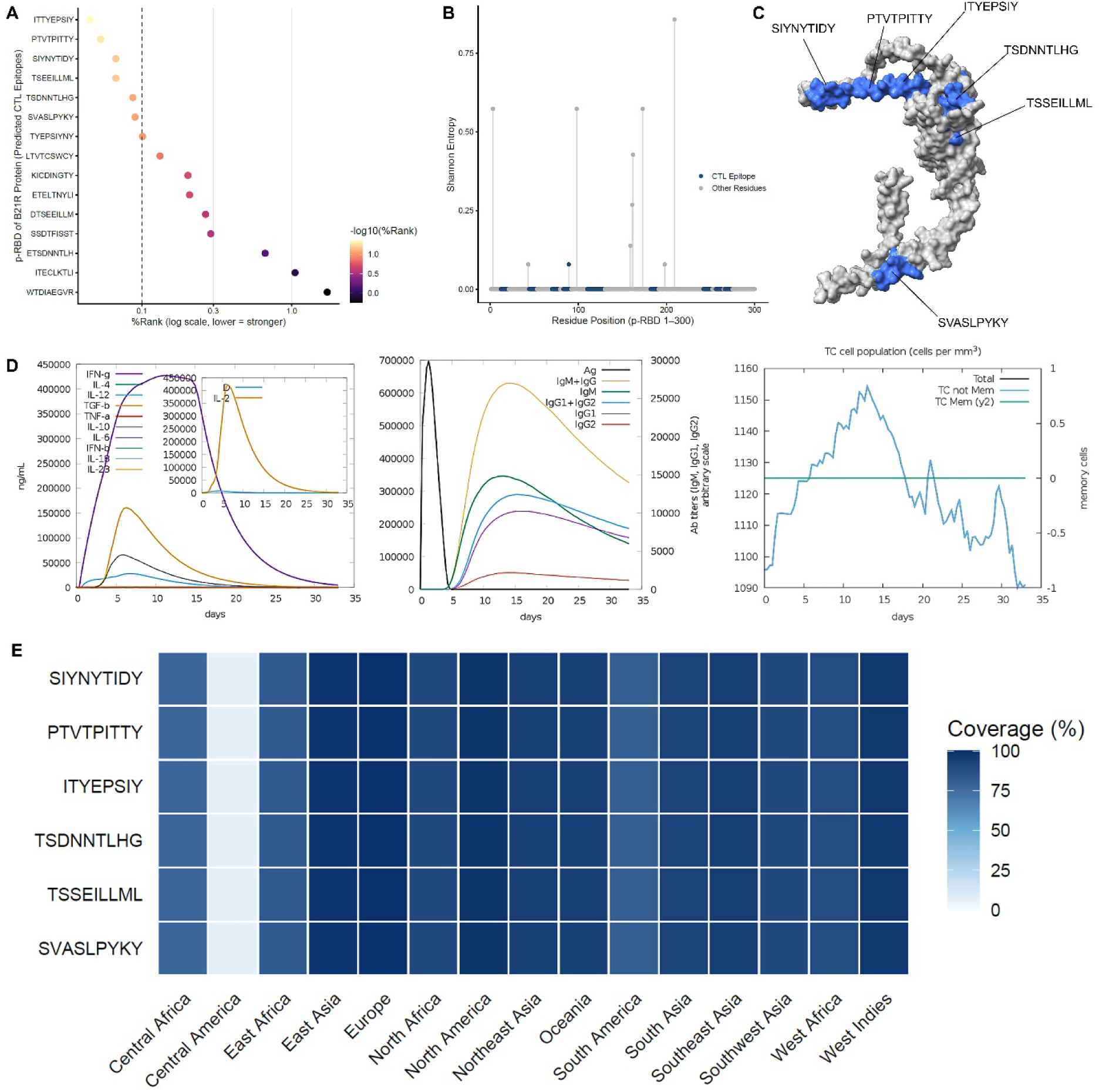
Identification and global population coverage of conserved CTL epitopes within the B21R p-RBD. **A.** Predicted MHC class I binding affinity of candidate CTL epitopes within the B21R p-RBD region based on PRIME analysis. Peptides are ranked according to %Rank (log scale), where lower values indicate stronger predicted binding and immunogenicity. Several epitopes, including ITTYEPSIY, PTVTPITTY, SIYNYTIDY, TSDNNTLHG, TSSEILLML, and SVASLPYKY, exhibited high immunogenic potential. **B.** Sequence conservation analysis across the B21R p-RBD (residues 1–300) showing Shannon entropy values at each residue position. Predicted CTL epitope residues (blue) display low entropy values, indicating strong conservation compared with more variable non-epitope regions (grey). **C.** Structural mapping of predicted CTL epitopes onto the AlphaFold-derived 3D structure of B21R p-RBD, illustrating that the epitopes are predominantly surface-exposed and spatially clustered within the putative receptor-binding region. **D.** C-ImmSim simulation showing host immune responses following antigen exposure showing rapid expansion of cytotoxic T cells and development of memory populations. Cytokine profiles indicate a Th1-biased immune response characterized by elevated IFN-γ and IL-2 production, accompanied by progressive antibody responses. **E.** Regional population coverage analysis showing predicted HLA class I coverage of selected CTL epitopes across major global populations. The selected peptides demonstrate high predicted population coverage (>80–99%) in most geographic regions, supporting their broad immunogenic potential.

C-ImmSim based agent-based immune simulation predicted successful p-RBD sequence antigen processing and presentation in specific HLA environments. The model showed a rapid increase in the number of cytotoxic CD8^+^ T cells on exposure to antigens by progressing into clonal proliferation followed by memory T-cell formation **(Figure 8C)**. There was a lack of minimal induction of anergic T-cell subsets. Cytokine profiling indicated a very strong Th1-biased response with high levels of both IFN-g and IL-2 which are in line with effective cellular antiviral immunity. Population coverage was analyzed by the HLA class I allele distribution which revealed a broad spectrum of global applicability of the selected set of CTL epitopes. Obtained pooled peptides had a projected coverage of over 90 per cent and highest coverage in Europe (99.42%), East Asia (98.08%), North American (97.91%), West Indies (96.86%), southeast Asia (94.45%) and Northeast Asia (94.36%). Another high coverage was also in African populations, where East (82.26%), West Africa (88.34%), North Africa (89.75%), and Central Africa (77.12) had the highest coverage, and South Asia (92.28) and Southwest Asia (89.53) had the highest coverage **(Figure 8D)**.

## Discussion

The global epidemiology of Mpox is imposing significant public health concern as the number of Mpox cases has significantly increased in the recent multinational outbreak [26]. Based on the number of confirmed cases on a 7 days average basis relative to the population per one million cases we have identified three clear distinct phases based on the different regions during 2022, 2023, 2024 and 2025 showing potential variation in viral transmission dynamics. We have identified that the 2022 outbreak of Mpox peaked in North America, South America, and Europe, and subsequently declined significantly in these regions by 2023, 2024 and 2025. However, in 2023, most of the cases of Mpox emerged from the Asia region with sporadic cases in Africa and Oceania. However, in the year 2024, the Mpox cases declined in Asia and later increased in 2025. Prominently, Mpox cases in the Africa region started in 2024 and peaked in 2025, making it an endemic hotspot [27].

In addition to Africa, Mpox cases were increased in Oceania in the Year 2024 with some cases in 2025. These findings suggest a transmission dynamic shift in Mpox epidemiology, possibly driven by genetic, immunological, and environmental factors [28, 29]. Importantly, when we tracked the Mpox cases from 2003, we have found that Mpox originated from Africa and subsequently transmitted to Europe and Asia which has further resulted in the current multinational outbreaks of Mpox which is similar to previous reports [8, 30].

To further understand the transmission dynamics of Mpox, we considered the role of B21R protein as one of the crucial structural proteins of MPXV which might be playing an important role in Mpox transmission [31], entry into the host target cells [32] and immune evasion strategies [22, 33]. As Mpox emerged in the year 1996, we performed our analysis using MPXV strains from the major outbreaks of Mpox and therefore we have retrieved various sequences from one geographical region and categorized them based on their collection date and geographical regions [34]. As we could not find much variation among the retrieved sequences from one geographical to other regions, we generated the normalized consensus sequences designated as year wise 2022, 2023, 2024 and 2025 MPXV strains for the current multinational outbreak of Mpox and compared with the previous major outbreak strains.

Our phylogenetic analysis revealed significant divergence of the recent MPXV strains from 1996 strain of MPXV where we found that MPXV strains from 2022, 2023, 2024 and 2025 are closely clustered. This data suggests an evolutionary trajectory that may have contributed to recent increase in Mpox transmission. This lineage divergence shows subtle mutations in B21R protein which may have played a role in viral adaptation and immune escape mechanism [35]. Upon detailed analysis of mutations across the strains, we have found 19 unique mutations in 2003 MPXV strain as compared with the reference Zaire-96-I-16 strain. Prominently, 17 of these mutations persisted in subsequent outbreaks whereas two were unique to the 2003 MPXV strain. Furthermore, the intermediate outbreak in 2017 introduced five novel mutations which were later retained in the 2022– 2025 Mpox outbreaks. In addition to these mutations, several novel mutations D209N, P722S, and M1741I, emerged in 2022, which were also present in 2023 and 2024 while 2025 strain exhibited only M1741I mutation whereas D209N and P722S mutations were absent. In addition to these mutations, the 2025 MPXV strain also shows two unique mutations T149D and D1877N. These gradual mutations and adaptation of the MPXV strain show an increase in viral fitness along with immune evasion strategies.

Importantly, we generated the structure of B21R protein using AlphaFold which showed higher confidence in intra-domain residue positioning. To investigate the role of B21R protein in Mpox evolution, we performed the solvent accessibility of the B21R protein and identified that the p-RBD and central ectodomain region exhibited the higher solvent accessibility, further strengthening the role of p-RBD region in contributing to its possible functional activity in receptor binding and immune recognition [36]. It is interesting to note that overlapping of the mutations on the residues L3, D98, T149, I173 and D209 on the AlphaFold-predicted structure indicated that they were concentrated in an uninterrupted patch of the surface and that they were not caused by random genetic drift. Conversely, those mutations that were located in buried sites or observed in other regions of the structure that were structurally constrained were less likely to affect the interactions with hosts. Taken together, these observations suggest that mutation exposed to the surface is a structurally interesting hot spot, which is typical of a p-RBD.

To further understand the role of mutations in B21R protein towards Mpox transmission, we determined the affinity of p-RBDs towards the CRD region of the DC-SIGN receptor, a critical host receptor for viral entry [23, 25]. Upon analyzing the above protein-protein interaction, we observed significant reduction of distance from 46.34Å in the Zaire-96-I-16 strain to 32.08Å in the 2025 strain of MPXV between pRBDs and CRD region of DC-SIGN receptor. This data is suggesting a progressive enhancement of receptor binding affinity and may be responsible for the increased MPXV transmission, supporting our hypothesis that evolution of MPXV has optimized host-receptor interactions to enhanced attachment and infection. Moreover, recent outbreak strains showed progressively more favorable binding free energies (ΔG) and reduced dissociation constants (KD).

Moreover, we performed mapping of N-linked and O-linked glycosylation sites across the analyzed MPXV strains. We have found that the N-linked glycans remain consistent across the MPXV strains whereas we observed transformation of O-linked glycans. We observed that N-linked sites were widely distributed across the p-RBD and therefore playing an important role in the protein stability and immune shielding. Whereas, the O-linked sites were found to be localized in the flexible loop region of p-RBD, representing its crucial role in the engagement of the CRD region of DC-SIGN. This pattern-specific remodeling of O-linked glycosylation motifs in the p-RBD indicates a possible antigenic diversification mechanism in which modifications in glycan positioning can potentially modify surface accessibility, presentation of epitopes and facilitate receptor interactions. This variability can be used in differentiating host interaction and immune recognition between recent MPXV strains. Nevertheless, the modulations of these O-linked glycosylation on functionality especially concerning the B-cell epitope concealment and the connection of host receptors are crucial. Therefore, we analyzed the antigenic divergences due to variation in O-linked glycosylation sites and we observed significant variation in linear B cell epitopes specifically at the p-RBDs across the analyzed strains which plays a crucial role in humoral immune escape mechanism, whereas the CTL epitopes remain consistent. These antigenic differences show that the recent MPXV strains evolved to escape the preexisting neutralizing antibody responses and therefore contributing to increased case numbers worldwide. Furthermore, the observed phylogenetic diversity of MPXV strains underscores the importance of designing a broad-spectrum vaccine candidate targeting conserved epitopes across different clades. Our data revealed crucial mutational hotspots of B21R protein and may be considered for the development of future Mpox vaccines focusing on emerging MPXV strains. By targeting the conserved B21R-CRD-DC-SIGN interacting residues and linear B cell epitopes, monoclonal antibody-based therapy can be developed to prevent the MPXV entry. Moreover, use of monoclonal antibodies recognizing glycosylated epitopes of B21R protein can improve the neutralization of emerging MPXV strains [37]. As the CTL epitopes remained consistent across the MPXV strains, development of a T cell-based vaccine using B21R protein is crucial to achieve [38]. Hence an ideal Mpox vaccine could reduce the risk of escape mutations and ensure lasting protection against future Mpox outbreaks.

Interestingly, p-RBD based CTL epitopes showed strong MHC-I binding affinity, TCR recognition scores. These CTL epitopes were found to be conserved across MPXV strains from 1996-2025. Notably, C-ImmSim based immune simulation anticipated a robust CD8^+^ T cell proliferation, stable T memory cell formation, and Th1 biased cytokine release, which included high levels of IFN-γ and IL-2, which are crucial for CD8^+^ T cell effector function and memory formation [39]. Moreover, the population coverage analysis showed that it is widely applicable in various HLA class I distributions by covering over 90 percent of various geographical locations. These results demonstrated that B21R p-RBD is a potential T cell-based vaccine candidate against MPXV as it has conserved and immunogenic CTL determinants that have the potential to induce a protective cellular response. However, its protective efficacy needs to be investigated in experimental setup.

In conclusion, our data provides novel insights into the molecular and epidemiological shift in the recent Mpox outbreaks. Our integrated global epidemiological data associated with structural and functional analysis revealed plausible MPXV global transmission. Based on mutational analysis of B21R protein across the MPXV strains and its affinity towards the CRD region of DC-SIGN receptor, we revealed plausible MPXV transmission and immune evasion strategy that can lead to development of potential therapeutics and effective T cell-based vaccine candidate based on p-RBD of B21R protein.

## Materials and Methods

### Global epidemiology of Mpox based on current outbreaks

To decipher the global epidemiological transformation in current outbreaks of Mpox, we analyzed the number of confirmed cases on a 7 days average basis relative to the population per one million cases using Our World in Data (https://ourworldindata.org/mpox) [40]. Visualization of spatial patterns of transmission was created by generating annual choropleth maps of Mpox incidence (cases per million) from 2022-2025 based on Natural Earth country boundary shapefiles accessed through the sf package in R. Epidemiological data were merged to spatial geometries based on standardized ISO3 country codes. All maps were plotted with geom_sf and a uniform color scale across years to enable one-to-one comparison of incidence intensity. Spatial projections were maintained in the WGS84 coordinate reference system (EPSG:4326). To stratify geographically by continent, six major geographic regions were selected as North America, South America, Europe, Asia, Africa and Oceania. Calculations of continental incidence were performed by aggregating country-level cases per million and presented as time-series plots between May 2022 and December 2025. Temporal trends were evaluated to identify the major outbreak waves and region-wise transmissions dynamics. On the basis of continent-wise temporal incidence curves, the dynamics of outbreaks were defined in certain epidemiological phases according to the appearance of the first global surge, regional persistence, reemergence in individual regions, and region-specific transmission patterns, in 2022, 2023, 2024, and 2025, respectively. The classification of the phases was determined according to the sustained rises or declines of the rolling incidence patterns. All data processing, statistical analysis, and visualization were performed in R version 4.3.1 using the packages tidyverse, zoo, ggplot2, sf, and rnaturalearth. To combine the epidemiology transformation with Mpox evolution, genomic sequences specifically for Clade II were obtained from Nextstrain [41]. Continent-wise distribution of sequences across years was performed to assess lineage expansion and geographic distribution. Phylodynamic trends were plotted using scatter plots of frequency of sequences vs sampling year.

### Phylogenetic analysis of MPXV and mutational mapping of B21R protein

To analyze the relationship among the emerging strains of MPXV, we have used OPG210 encoding protein B21R, which is one of the surface membrane glycoproteins and exhibits high immunogenicity. We first derived the sequences of B21R protein from the year 1996, 2003, 2017 and continent wise consensus sequences from year 2022, 2023, 2024 and 2025 from Monkeypox Virus Resource (MPoxVR) version 1.0 [34]. As the current epidemiology leverages us to use more than one MPXV strain per continent, we retrieved the MPXV B21R protein sequences from all continents to understand the Mpox evolution in the current outbreaks and continent wise phylogenetics were performed using MEGA software [42]. Subsequently, EMBOSS Cons was used to generate the year wise consensus sequence [43]. To identify the crucial mutations in the current MPXV strains, we performed the multiple sequence alignment of MPXV strains from 1996, 2003, 2017 using Clustal Omega [44]. The mutations were compared with the reference strain of MPXV Zaire 1996. Hierarchical clustering based on the Pearson correlation similarity across binary mutation vectors was used to visualize the temporal mutation patterns across the B21R protein among the MPXV strains. The agglomerative clustering was performed using the linkage algorithm and the dissimilarity was computed as 1−r. Heatmap was generated using the ComplexHeatmap framework in R and chronological order of the sampling years was maintained [45]. Consequently, we performed the phylogenetic analysis based on the above normalized sequences and the phylogenetic tree was generated using TreeViewer [46].

### Ab initio structure modeling of B21R protein and solvent accessibility analysis

As the structure of the B21R protein is not yet deciphered we generated its tertiary model using AlphaFold server BETA through *ab initio* protein modelling as the template for the B21R protein is not available [47]. AlphaFold uses its trained neural networks to predict inter-residue distances and orientations, which are crucial for building accurate structures even without evolutionary guidance. The PAE matrix were generated based on the output file which represents the confidence of pairwise residue alignment with darker diagonal positions indicating greater confidence in the positioning of residues within one domain and lighter off-diagonal positions indicating flexibility of different domains. To decipher the potential domains involved in the interaction with the host receptor, we performed the Solvent-accessible surface area of B21R protein in UCSF ChimeraX [48]. The residues were clustered into structural domains (N-terminal, central ectodomain, C-terminal domain, and TM/tail) according to the number of the residues. The distribution of region-wise solvent accessibility was plotted in R using geom tile of the ggplot2 package to plot a heatmap of residues in domains. Categorical bars of annotation UniProt domains were superimposed on the structural context as categorical bars below the residue axis. To compare the solvent accessibility of structural domains, relative SASA values were plotted in ggplot2 with boxplot representation. The statistical test of the differences between the domains was performed by the non-parametric Kruskal-Wallis test. To visualize the solvent-exposed critical residues and common mutations spatial localization at the p-RBD, selected mutational hotspots were overlaid on the three-dimensional structure in ChimeraX.

### Protein–protein interaction analysis of B21R–DC-SIGN complexes

AlphaFold2 pipeline was used to generate three-dimensional models of p-RBDs of B21R protein for each Mpox-associated outbreak as 1996, 2003, 2017, 2022, 2023 and 2025. The crystal structure of the carbohydrate recognition domain (CRD) of DC-SIGN was obtained in the protein data bank (PDB ID:1SL4) [49]. Calcium-binding sites such as E347, N349, E354, N365 and D366 were used for the protein-protein docking of p-RBDs and CRD region of DC-SIGN using HADDOCK 2.4 [50]. HADDOCK score (combination of van der Waals, electrostatic, desolvation and restraint violation energies) was used to select the top-ranked clusters. The most ideal cluster in terms of the lowest HADDOCK score and good interface properties was chosen to further the analysis of each outbreak strain. Intermolecular interaction in each B21R-DC-SIGN complex was predicted on the MAPIYA server [51]. The level of interaction at the residue level was done to identify the interface contacts, such as hydrogen bonds, van der Waals forces, polar/apolar contributions. Geometric complementarity between the two proteins was estimated using contact distances.

### Variation of O-linked glycosylation in B21R proteins of MPXV

As the B21R protein is a glycoprotein which exhibits affinity towards the carbohydrate recognition domain of DC-SIGN receptor, deciphering its glycosylation sites is crucial to understand the transmission and virulence of the recent MPXV strains. Using artificial neural networks, we determined the variations in N- and O-linked glycans [52, 53]. A compiled and visualized glycosylation profile, in terms of conservation strains and variation strains, was produced using R (ggplot2), and allowed an evaluation of conservation and variation in the putative receptor-binding domain (p-RBD). To assess the mechanical effects of glycosylation, these sites were overlaid to the modeled three-dimensional structure of B21R and representative proxy glycans were computationally conjugated to individual residues. Due to the absence of explicit heparan sulfate disaccharide templates for direct protein attachment in GlycoShape, O-linked glycosylation was modeled using a minimal O-glycan proxy to approximate the steric and electrostatic contribution of HS-like glycans [54]. Basic clash resolution and rotamer adjustment were used to get the glycan conformations to a realistic spatial orientation. Visualization of the final glycosylated models was used to analyze spatial distribution, surface exposure and possible overlap with predicted B-cell epitopes in order to give structural sense to glycosylation conservation and variation between MPXV strains.

### Antigenic variation of B21R protein

To understand the antigenic variation of the B21R protein we determined the variations in B cell epitopes and CTL epitopes among the MPXV strains Zaire-96, 2003, 2017, 2022, 2023, 2024, and 2025. The B cell epitopes were determined by the antibody epitope prediction [55] and CTL epitopes were determined by the T cell epitope prediction tools [56], and only high-confidence epitopes above the defined scoring threshold were considered for downstream analysis The peptide sequences of each strain, prediction scores, epitope start, and end positions were tabulated. Overlapping epitopes consolidated into canonical antigenic regions to enable a consistent cross strain comparison. Diversity was evaluated by the analysis of positional changes, change in lengths, substitutions in sequences and alterations in scores of epitopes prediction within different strains. As part of making comparative visualizations, epitope scores in specific antigenic regions were plotted as heatmaps in R with ggplot2, with color intensities indicative of relative antigenicity. To measure surface exposure and spatial clustering of the selected epitope regions, structural mapping of the regions on the three-dimensional model of B21R was performed.

### B21R p-RBD–induced cellular immunity for the potential T cell-based vaccine candidate

To strengthen the rationale of using p-RBD as a T cell-based vaccine for Mpox, we determined the T-cell immunogenic potential of the p-RBD protein domain of the MPXV B21R (OPG210) protein using an integrated immunoinformatics methodology. Prediction of CD8^+^ T cell epitopes was done with the PRIME v2.1 which is a mix of the MixMHCpred based antigen presentation modeling and TCR recognition scoring [57]. The 9-mer peptide repertoire of the p-RBD sequence was screened against typical alleles of the HLA class I (HLA-A01:01, HLA-A25:01, HLA-B07:02 and HLA-B18:01). The peptides were ranked according to the values of the percentile rank of best allele and lower percentile rank values were associated with predicted immunogenicity. To determine the stability of evolution, conservancy analysis of the predicted epitopes was performed between the MPXV strains (1996-2025). Subsequently, to assess functions of immune dynamics, an agent-based immune simulation using the C-ImmSim platform was performed with a model of vaccination on the entire p-RBD sequence [58]. The results were simulated using 100-time steps with single antigen injection and adjuvant support in specified HLA conditions (HLA-A01:01, HLA-B07:02 and HLA-DRB1:01). The model simulated the antigen processing, the presentation of MHC class 1 and class II, the expansion of CD4^+^ and the CD8^+^ T cells, cytokines and the formation of memory cells. Predicted cellular immune responses were measured by analyzing output parameters such as the expansion of cytotoxic T-cells, Th polarization, and cytokine profiles. The IEDB Population Coverage tool was used to compare population coverage of the predicted CTL epitopes (http://tools.iedb.org/population/) [59]. HLA class I alleles that were analyzed were as follows: HLA-A01:01, HLA-A02:01, HLA-A02:03, HLA-A02:06, HLA-A03:01, HLA-A11:01, HLA-A24:02, HLA-A26:01, HLA-A30:01, HLA-A30:02, HLA-A31:01, HLA-A32:01, HLA-A33:01, HLA-B07:02, HLA-B08:01, HLA-B15:01, HLA-B27:05, HLA-B35:01, HLA-B40:01, HLA-B44:02, HLA-B44:03, HLA-B51:01, HLA-B53:01, HLA-B57:01, and HLA-B*58:01. Population coverage was analyzed based on 15 different regions of the globe by calculating the HLA genotypic frequencies and epitope-HLA binding combinations. The population coverage was projected as a percentage and it was analyzed by heatmap in R.

## ADDITIONAL INFORMARTION

The online version contains supplementary material available.

## ACKNOWLEDGEMENTS

The authors are grateful to the Vice Chancellor, King George’s Medical University (KGMU) Lucknow, for the encouragement for this work. The authors have no other relevant affiliations or financial involvement with any organization or entity with a financial interest in or financial conflict with the subject matter or materials discussed in the manuscript apart from those disclosed.

## AUTHOR CONTRIBUTIONS

SK^1^ and SKS conceived the idea and planned the study and contributed equally to this work as a first author. SK and ASH collected the data, devised the initial draft, and reviewed the final draft. SK^1^, ASH, SK^2^, JTP, ASA and SKS finalized the draft for submission. All authors read and approved the final version of the manuscript.

