## Supplementary File for "Adaptive Remodeling of the MPXV B21R Receptor-Binding Domain Enhances DC-SIGN Interaction and Identifies Conserved CTL Targets for T-Cell Vaccine Development"

**Prof. Shailendra K. Saxena**

Centre for Advanced Research (CFAR), Faculty of Medicine,

King George’s Medical University (KGMU), Lucknow 226003, India

**Table S1.** Accession IDs of B21R protein sequences of MPXV used in the study from 1996-2025.

| **1996** | | | | | | |
| --- | --- | --- | --- | --- | --- | --- |
| OPG210 | NP_536609.1 | | | | | |
| **2003** | | | | | | |
| OPG210 | DQ011157.1 | | | | | |
| **2017** | | | | | | |
| OPG210 | UXL78637.1, UXL77766.1, UXL78459.1 | | | | | |
| **Jan 2022–Dec 2022** | | | | | | |
| OPG210 | **North America** | **South America** | **Europe** | **Africa** | **Asia** | **Oceania** |
|  | NY-NYCPHL-001204 | ANT-LDSP-ANT-MPX-28354C | hMpxV/Germany/BE-ChVir28607/2022 | TRM328 | MPXV-ROK-P002-2022 | 43_A |
|  | MPXV_USA_2022_NY0048 | LIM-INS008 | MPXV/Germany/2022/ON/RKI932 | TRM306 | MPXV/human/Japan/Tokyo/2022/TKY220165 | 7_A |
|  | MPXV_USA_2022_TX0054 | LIM-INS-021 | Monkeypox/PT0046/2022 | TRM294 | hMpxV/Philippines/RITM-002/2022 | 54_A |
|  | MPXV_USA_2022_CA0004 | LIM-INS-003 | MPX_K1dQNt_9000223 | TRM311 | MPXV-ROK-P004-2022 | 4_A |
|  | MPXV_USA_2022_PA0003 | LIM-INS-018 | MPX_BhVKjm_9000232 | TRM303 | MPXV-ROK-P1-2022 | 42_C |
|  | MPXV_USA_2022_MI0012 | LIM-INS-015 | MPXV/UZ_REGA_10 | TRM081 | MPXV/human/Japan/Tokyo/TKY220091/2022 | 38_A |
| **Jan 2023–Dec 2023** | | | | | | |
| OPG210 | **North America** | **South America** | **Europe** | **Africa** | **Asia** | **Oceania** |
|  | hMpxV/USA/CA-CDPH-1M1000558/2023 | - | MPXV/Germany/2023/ON/RKI1077 | VSP205 | MPXV-ROK-P53/03-2023 | - |
|  | hMpxV/USA/CA-CDPH-1M1000424/2023 | - | MPXV/Germany/2023/ON/RKI1057 | VSP212 | MPXV-ROK-P53/05-2023 | - |
|  | hMpxV/USA/CA-CDPH-1M1000427/2023 | - | MPXV/Germany/2023/ON/RKI1054 | RDC-NKV-GOM-MPOX-010 | MPXV/human/Japan/Tokyo/2023/TKY220343 | - |
|  | hMpxV/USA/CA-CDPH-1M1000449/2023 | - | MPXV/PT0810/2023 | VSP192 | MPXV/human/Japan/Tokyo/2023/TKY220318 | - |
|  | hMpxV/USA/CA-CDPH-1M1000426/2023 | - | MPXV/PT0809/2023 | 24MPX0009C | MPXV/human/Japan/Tokyo/2023/TKY220310 | - |
|  | NY-NYCPHL-001187 | - | 02022023_gluteal | VSP190 | hMpxV/THA/V241-0052/2023 | - |
| **Jan 2024–Dec 2024** | | | | | | |
| OPG210 | **North America** | **South America** | **Europe** | **Africa** | **Asia** | **Oceania** |
|  | NYCPHL-0001368 | - | MPXV/Germany/2024/ON/RKI1101 | 24MPX0038V | MPXV/human/Japan/Tokyo/NCGM240303/2024 | hMPXV/Australia/VIC-7288/2024 |
|  | NYCPHL-0001364 | - | MPXV/Germany/2024/ON/RKI1103 | 24MPX0164V | CU-ID24110 | hMPXV/Australia/VIC-8550/2024 |
|  | Monkeypox virus/Human/USA/CA-LACPHL-MA00662/2024 | - | Monkeypox/PT0829/2024 | 24MPX0194V | - | hMPXV/Australia/VIC-6919/2024 |
|  | NYCPHL-0001365 | - | MPXV/PT0822/2024 | 24MPX0209V | - | hMPXV/Australia/VIC-6170/2024 |
|  | NYCPHL-0001367 | - | MPXV/Germany/2024/ON/RKI1107 | 24MPX0240V | - | hMPXV/Australia/VIC-5972/2024 hMPXV/Australia/VIC-9145/2024 |
|  | - | - | MPXV/Germany/2024/ON/RKI1100 | 24MPX0220V | - | hMPXV/Australia/VIC-7288/2024 |
| **Jan 2025–Dec 2025** | | | | | | |
| OPG210 | **North America** | **South America** | **Europe** | **Africa** | **Asia** | **Oceania** |
|  | [MPXV_USA_2025_NH0100](https://www.ncbi.nlm.nih.gov/nuccore/PV294984.1) | - | [MPXV/Germany/2025/ON/RKI1264](https://www.ncbi.nlm.nih.gov/nuccore/PV186878.1) | [MpxvSL02](https://www.ncbi.nlm.nih.gov/nuccore/PV683567.1) | [hMpxV/Thailand/DDCBIDI-001/2025](https://www.ncbi.nlm.nih.gov/nuccore/PQ963869.1) | - |
|  | [WA-UW-040077](https://www.ncbi.nlm.nih.gov/nuccore/PV031938.1) | - | [MPXV/Germany/2025/ON/RKI1347](https://www.ncbi.nlm.nih.gov/nuccore/PV643753.1) | [MpxvSL01](https://www.ncbi.nlm.nih.gov/nuccore/PV683566.1) | - | - |
|  | [WA-UW-97972](https://www.ncbi.nlm.nih.gov/nuccore/PV592658.1) | - | [MPXV/Germany/2025/ON/RKI1369](https://www.ncbi.nlm.nih.gov/nuccore/PV683321.1) | - | - | - |
|  | [WA-UW-89335](https://www.ncbi.nlm.nih.gov/nuccore/PV697338.1) | - | [MPXV/Germany/2025/ON/RKI1351](https://www.ncbi.nlm.nih.gov/nuccore/PV643757.1) | - | - | - |
|  | [WA-UW-52066](https://www.ncbi.nlm.nih.gov/nuccore/PV697337.1) | - | [MPXV/Germany/2025/ON/RKI1363](https://www.ncbi.nlm.nih.gov/nuccore/PV643769.1) | - | - | - |
|  | [MPXV_USA_2025_NY0100](https://www.ncbi.nlm.nih.gov/nuccore/PV448277.1) | - | [MPXV/Germany/2025/ON/RKI1364](https://www.ncbi.nlm.nih.gov/nuccore/PV643770.1) | - | - | - |

*^§^To normalize the sequencing error, depending on the availability a maximum of 6 MPXV sequences from each geographical region were retrieved to represent the multinational Mpox outbreaks.*

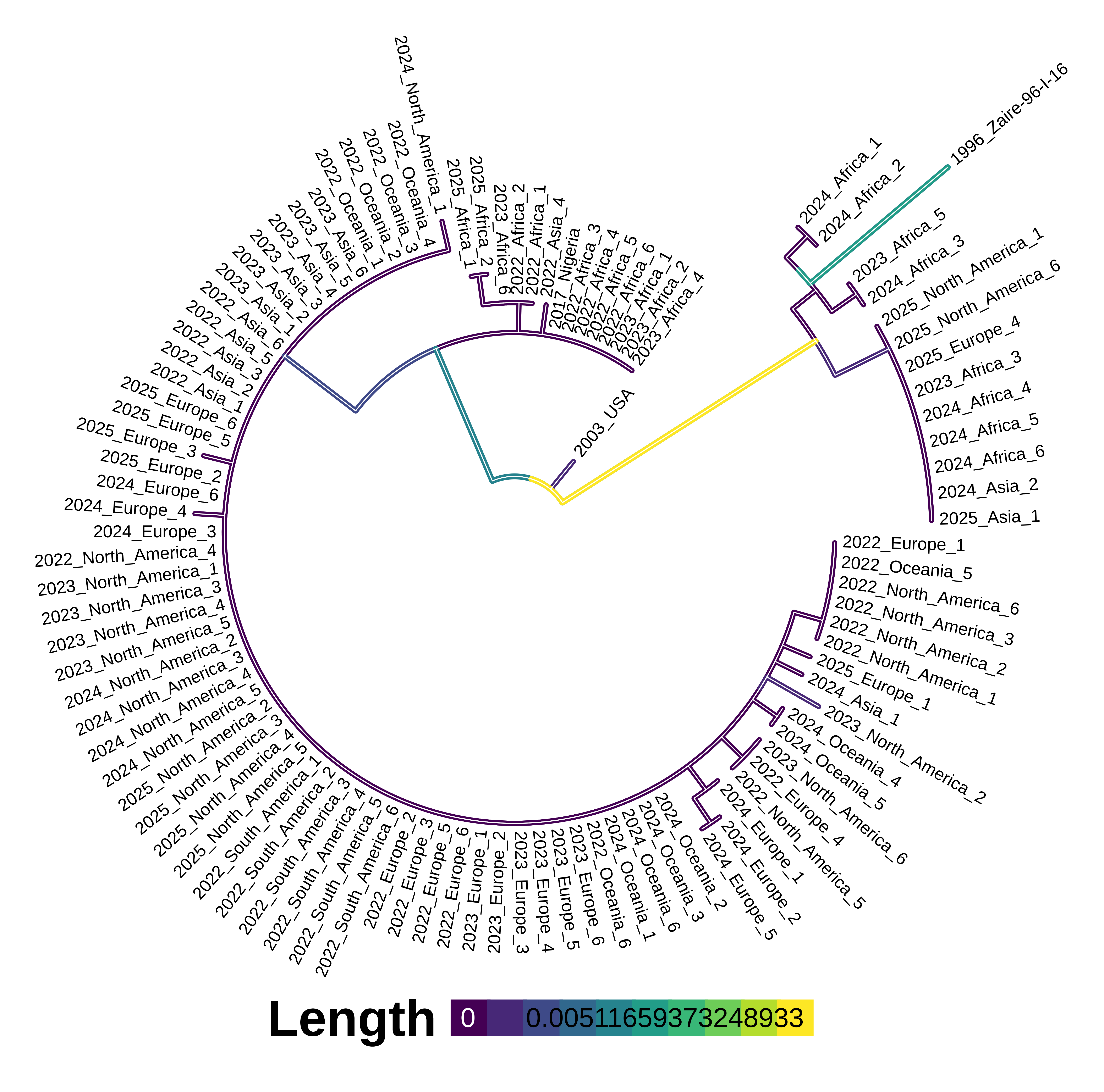

Figure S1. Phylogenetic analysis of the recent multinational outbreak related continent wise MPXV strains.

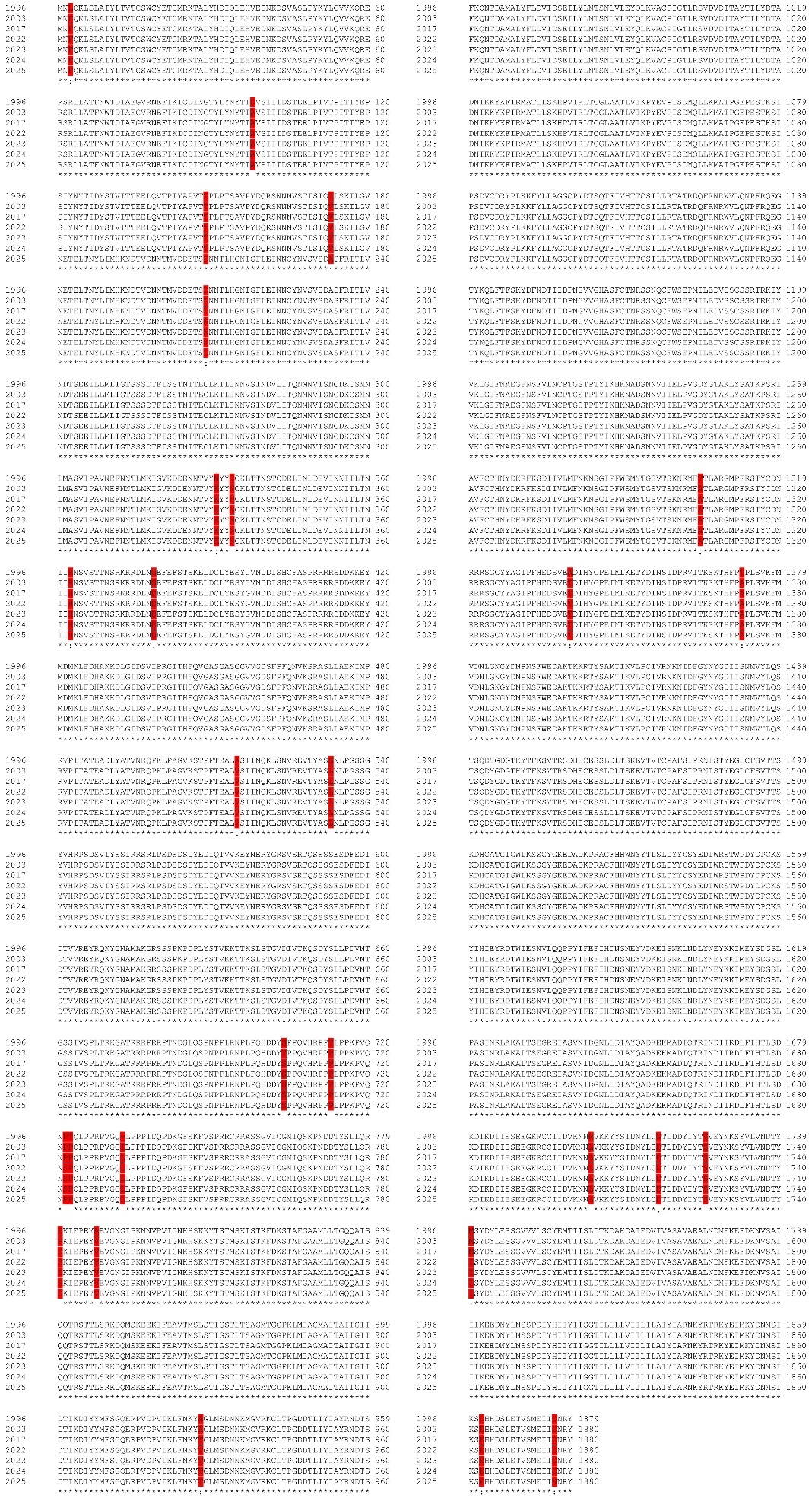

Figure S2. Multiple Sequence alignment based on the year wise consensus MPXV strains. Red color highlights the mutational hotspots in the B21R protein.

Table S2. Contact map residues and distance between B21R and CRD region of DC-SIGN receptor across the MPXV strain from 1996 to 2025.

| B21R | CRD-Region | Distance (A) | B21R | CRD-Region | Distance (A) | B21R | CRD-Region | Distance (A) | B21R | CRD-Region | Distance (A) | B21R | CRD-Region | Distance (A) | B21R | CRD-Region | Distance (A) | B21R | CRD-Region | Distance (A) |
| --- | --- | --- | --- | --- | --- | --- | --- | --- | --- | --- | --- | --- | --- | --- | --- | --- | --- | --- | --- | --- |
| Zaire-96-I-16-B21R | | | **2003-MPXV-B21R** | | | **2017-MPXV-B21R** | | | **2022-MPXV-B21R** | | | **2023-MPXV-B21R** | | | **2024-MPXV-B21R** | | | **2025-MPXV-B21R** | | |
| T96 | **E347** | 4.918 | T96 | **E347** | 7.697 | T96 | **E347** | 7.383 | Y95 | **E347** | 6.836 | A98 | **E347** | 4.499 | Y95 | **E347** | 7.923 | T96 | **E347** | 4.918 |
| I97 |  | 6.184 | I97 |  | 6.022 | I97 |  | 6.285 | T96 |  | 3.124 | V99 |  | 4.945 | T96 |  | 3.446 | I97 |  | 6.184 |
| D98 |  | 3.047 | A98 |  | 3.633 | A98 |  | 3.102 | I97 |  | 6.239 | S100 |  | 3.196 | I97 |  | 4.486 | D98 |  | 3.047 |
| V99 |  | 5.779 | V99 |  | 3.201 | V99 |  | 4.535 | A98 |  | 6.796 | I101 |  | 6.62 | A98 |  | 3.333 | V99 |  | 5.779 |
| S100 |  | 5.63 | S100 |  | 3.743 | S100 |  | 3.982 | H214 |  | 6.362 | I102 |  | 7.551 | V99 |  | 6.04 | S100 |  | 5.63 |
| H214 |  | 6.94 | I101 |  | 6.538 | I101 |  | 7.476 | D232 |  | 6.698 | R236 |  | 6.659 | S100 |  | 7.052 | H214 |  | 6.94 |
| S234 |  | 6.807 | H214 |  | 7.877 | H214 |  | 7.079 | S234 |  | 5.356 | E246 |  | 6.999 | H214 |  | 7.515 | S234 |  | 6.807 |
| R236 |  | 7.04 | S234 |  | 7.807 | S234 |  | 5.832 | F235 |  | 7.381 | I247 |  | 5.84 | S234 |  | 6.552 | R236 |  | 7.04 |
| L248 |  | 5.737 | R236 |  | 5.333 | F235 |  | 7.143 | R236 |  | 7.683 | L248 |  | 3.49 | R236 |  | 6.145 | L248 |  | 5.737 |
| L249 |  | 5.142 | E246 |  | 5.867 | R236 |  | 4.726 | L248 |  | 5.883 | L249 |  | 5.714 | L248 |  | 4.68 | L249 |  | 5.142 |
| M250 |  | 3.891 | I247 |  | 6.301 | I237 |  | 7.688 | L249 |  | 7.028 | M250 |  | 3.372 | L249 |  | 6.245 | M250 |  | 3.891 |
| L251 |  | 7.74 | L248 |  | 3.25 | T238 |  | 7.212 | M250 |  | 3.502 | Q286 |  | 7.111 | M250 |  | 3.504 | L251 |  | 7.74 |
| N289 |  | 5.539 | L249 |  | 4.63 | E246 |  | 5.19 | L251 |  | 5.382 | N287 |  | 2.758 | L251 |  | 5.595 | N289 |  | 5.539 |
| V290 |  | 7.407 | M250 |  | 3.509 | I247 |  | 6.884 | T252 |  | 2.788 | M288 |  | 5.314 | T252 |  | 6.236 | V290 |  | 7.407 |
| T291 |  | 4.266 | L251 |  | 7.447 | L248 |  | 3.188 | G253 |  | 6.05 | N289 |  | 5.245 | M288 |  | 7.412 | T291 |  | 4.266 |
| T96 | **N349** | 7.993 | N287 |  | 5.558 | L249 |  | 5.656 | N289 |  | 7.525 | I97 | **N349** | 7.714 | N289 |  | 3.201 | T96 | **N349** | 7.993 |
| I97 |  | 7.387 | M288 |  | 6.524 | M250 |  | 3.798 | V290 |  | 7.638 | A98 |  | 6.247 | V290 |  | 6.049 | I97 |  | 7.387 |
| D98 |  | 7.298 | N289 |  | 6.471 | L251 |  | 7.98 | T291 |  | 3.925 | V99 |  | 7.822 | T291 |  | 4.332 | D98 |  | 7.298 |
| V99 |  | 5.93 | T212 | **N349** | 7.328 | N287 |  | 5.708 | S292 |  | 6.748 | H214 |  | 6.3 | T96 | **N349** | 7.492 | V99 |  | 5.93 |
| S100 |  | 6.704 | L213 |  | 6.1 | M288 |  | 7.165 | N293 |  | 5.603 | G215 |  | 6.431 | A98 |  | 7.975 | S100 |  | 6.704 |
| L213 |  | 6.94 | H214 |  | 3.544 | N289 |  | 4.665 | K296 |  | 7.159 | N216 |  | 7.542 | V99 |  | 6.577 | L213 |  | 6.94 |
| H214 |  | 3.12 | G215 |  | 3.666 | V290 |  | 7.212 | Y188 | **N349** | 7.039 | D232 |  | 7.441 | L213 |  | 6.213 | H214 |  | 3.12 |
| G215 |  | 4.198 | N216 |  | 5.606 | T291 |  | 6.721 | H214 |  | 3.358 | A233 |  | 6.692 | H214 |  | 3.235 | G215 |  | 4.198 |
| N216 |  | 6.355 | D232 |  | 7.188 | Y188 | **N349** | 7.995 | G215 |  | 3.111 | S234 |  | 3.247 | G215 |  | 4.395 | N216 |  | 6.355 |
| I217 |  | 7.874 | A233 |  | 7.508 | T212 |  | 5.631 | N216 |  | 3.29 | F235 |  | 4.903 | N216 |  | 6.549 | I217 |  | 7.874 |
| D232 |  | 7.54 | S234 |  | 4.264 | L213 |  | 3.035 | I217 |  | 5.16 | R236 |  | 5.602 | D232 |  | 7.584 | D232 |  | 7.54 |
| A233 |  | 6.136 | F235 |  | 2.997 | H214 |  | 3.197 | G218 |  | 5.97 | L248 |  | 3.47 | A233 |  | 6.647 | A233 |  | 6.136 |
| S234 |  | 3.007 | R236 |  | 4.632 | G215 |  | 3.678 | D232 |  | 2.999 | L249 |  | 5.01 | S234 |  | 3.265 | S234 |  | 3.007 |
| F235 |  | 2.927 | I237 |  | 7.208 | N216 |  | 5.354 | A233 |  | 4.931 | M250 |  | 3.167 | F235 |  | 3.196 | F235 |  | 2.927 |
| R236 |  | 3.804 | L248 |  | 3.823 | I217 |  | 7.856 | S234 |  | 3.27 | L251 |  | 6.529 | R236 |  | 3.072 | R236 |  | 3.804 |
| I237 |  | 7.018 | M250 |  | 5.091 | D232 |  | 7.562 | F235 |  | 4.942 | T252 |  | 6.911 | I237 |  | 6.195 | I237 |  | 7.018 |
| I247 |  | 6.92 | T252 |  | 7.326 | A233 |  | 6.512 | R236 |  | 7.915 | T96 | **E354** | 4.723 | E246 |  | 7.385 | I247 |  | 6.92 |
| L248 |  | 3.052 | T15 | **E354** | 7.929 | S234 |  | 2.709 | M250 |  | 6.686 | I97 |  | 7.338 | I247 |  | 6.818 | L248 |  | 3.052 |
| L249 |  | 3.773 | W18 |  | 5.837 | F235 |  | 2.879 | T252 |  | 5.76 | A98 |  | 7.204 | L248 |  | 3.439 | L249 |  | 3.773 |
| M250 |  | 3.601 | C19 |  | 6.79 | R236 |  | 3.236 | W18 | **E354** | 6.628 | M250 |  | 6.046 | L249 |  | 4.363 | M250 |  | 3.601 |
| L251 |  | 6.602 | T22 |  | 5.052 | I237 |  | 5.095 | C19 |  | 7.27 | L251 |  | 7.526 | M250 |  | 3.522 | L251 |  | 6.602 |
| T252 |  | 7.7072 | Y95 |  | 7.762 | T238 |  | 7.4 | E21 |  | 7.914 | T252 |  | 7.069 | L251 |  | 6.748 | T252 |  | 7.072 |
| N216 | **E354** | 7.675 | T96 |  | 6.188 | E246 |  | 7.707 | T22 |  | 4.786 | N289 |  | 6.125 | T252 |  | 6.111 | N216 | **E354** | 7.675 |
| D232 |  | 6.88 | D232 |  | 7.145 | L248 |  | 6.285 | T252 |  | 7.57 | V290 |  | 7.718 | S292 |  | 3.349 | D232 |  | 6.88 |
| S234 |  | 7.271 | M250 |  | 7.356 | L249 |  | 7.51 | K296 |  | 5.922 | T291 |  | 4.123 | T22 | **E354** | 6.732 | S234 |  | 7.271 |
| M250 |  | 5.768 | T252 |  | 4.783 | M250 |  | 5.028 | T22 | **N365** | 7.865 | S292 |  | 5.884 | T96 |  | 7.409 | M250 |  | 5.768 |
| T252 |  | 7.105 | G253 |  | 7.559 | T252 |  | 7.488 | T96 |  | 5.753 | N293 |  | 5.145 | D232 |  | 5.444 | T252 |  | 7.105 |
| T96 | **N365** | 5.863 | N293 |  | 6.057 | C19 | **E354** | 6.376 | M250 |  | 6.721 | K296 |  | 6.848 | S234 |  | 7.724 | D98 | **N365** | 6.169 |
| D98 |  | 6.169 | K296 |  | 2.629 | Y20 |  | 6.46 | T252 |  | 6.964 | A98 | **N365** | 7.631 | M250 |  | 5.266 | M250 |  | 6.846 |
| M250 |  | 6.846 | T96 | **N365** | 5.875 | T96 |  | 6.257 | T291 |  | 5.126 | N287 |  | 6.554 | T252 |  | 5.993 | N289 |  | 6.772 |
| N289 |  | 6.772 | I97 |  | 7.876 | N216 |  | 7.461 | S292 |  | 5.204 | N289 |  | 4.493 | N293 |  | 7.805 | V290 |  | 7.745 |
| V290 |  | 7.745 | A98 |  | 6.355 | D232 |  | 5.141 | N293 |  | 4.621 | T291 |  | 7.442 | K296 |  | 5.277 | T291 |  | 3.939 |
| T291 |  | 3.939 | M250 |  | 6.099 | S234 |  | 6.57 | K296 |  | 3.593 | T96 | **D366** | 7.101 | T96 | **N365** | 5.793 | S292 |  | 6.592 |
| S292 |  | 6.592 | N289 |  | 5.753 | T243 |  | 7.354 | C19 | **D366** | 7.945 | I97 |  | 7.571 | A98 |  | 7.533 | N293 |  | 5.629 |
| N293 |  | 5.629 | V290 |  | 6.015 | M250 |  | 7.076 | T22 |  | 3.763 | A98 |  | 4.811 | M250 |  | 5.782 | K296 |  | 4.265 |
| K296 |  | 4.265 | T291 |  | 3.172 | T252 |  | 5.11 | C23 |  | 7.965 | M250 |  | 5.534 | N289 |  | 5.475 | N94 | **D366** | 7.452 |
| N94 | **D366** | 7.452 | S292 |  | 6.436 | T291 |  | 7.856 | T96 |  | 5.696 | N287 |  | 7.635 | V290 |  | 6.894 | Y95 |  | 7.103 |
| Y95 |  | 7.103 | N293 |  | 6.256 | N293 |  | 6.421 | M250 |  | 7.805 | M288 |  | 7.322 | T291 |  | 3.367 | T96 |  | 3.638 |
| T96 |  | 3.638 | K296 |  | 7.264 | K296 |  | 5.498 | T252 |  | 5.176 | N289 |  | 2.872 | S292 |  | 5.303 | I97 |  | 7.748 |
| I97 |  | 7.748 | T96 | **D366** | 3.901 | T96 | **N365** | 5.978 | G253 |  | 7.726 | V290 |  | 6.359 | N293 |  | 5.121 | N98 |  | 6.685 |
| D98 |  | 6.685 | I97 |  | 7.078 | I97 |  | 6.311 | T291 |  | 6.974 | T291 |  | 3.661 | D295 |  | 7.843 | M250 |  | 5.566 |
| M250 |  | 5.566 | A98 |  | 6.976 | A98 |  | 3.891 | S292 |  | 6.294 | S292 |  | 7.819 | K296 |  | 5.057 | L251 |  | 7.988 |
| L251 |  | 7.988 | D232 |  | 7.569 | V99 |  | 7.299 | N293 |  | 4.026 | - | - | - | N94 | **D366** | 6.76 | T252 |  | 4.572 |
| T252 |  | 4.572 | S234 |  | 7.912 | S100 |  | 7.762 | K296 |  | 2.642 | - | - | - | Y95 |  | 6.506 | G253 |  | 7.865 |
| G253 |  | 7.865 | M250 |  | 3.161 | L248 |  | 7.52 | - | - | - | - | - | - | T96 |  | 2.876 | T291 |  | 4.051 |
| T291 |  | 4.051 | L251 |  | 7.524 | M250 |  | 3.323 | - | - | - | - | - | - | I97 |  | 7.214 | S292 |  | 6.81 |
| S292 |  | 6.81 | T252 |  | 4.692 | N287 |  | 7.896 | - | - | - | - | - | - | D232 |  | 7.528 | N293 |  | 4.445 |
| N293 |  | 4.445 | G253 |  | 7.677 | M288 |  | 7.635 | - | - | - | - | - | - | M250 |  | 3.984 | L296 |  | 3.438 |
| K296 |  | 3.438 | T291 |  | 5.933 | N289 |  | 3.284 | - | - | - | - | - | - | L251 |  | 6.471 | - | - | - |
| - | - | - | N293 |  | 6.422 | V290 |  | 5.963 | - | - | - | - | - | - | T252 |  | 3.746 | - | - | - |
| - | - | - | K296 |  | 4.648 | T291 |  | 3.201 | - | - | - | - | - | - | G253 |  | 6.41 | - | - | - |
| - | - | - | - | - | - | S292 |  | 7.479 | - | - | - | - | - | - | N289 |  | 7.385 | - | - | - |
| - | - | - | - | - | - | T96 | **D366** | 4.906 | - | - | - | - | - | - | V290 |  | 7.693 | - | - | - |
| - | - | - | - | - | - | I97 |  | 7.851 | - | - | - | - | - | - | T291 |  | 3.735 | - | - | - |
| - | - | - | - | - | - | A98 |  | 6.258 | - | - | - | - | - | - | S292 |  | 5.814 | - | - | - |
| - | - | - | - | - | - | S234 |  | 7.566 | - | - | - | - | - | - | N293 |  | 3.528 | - | - | - |
| - | - | - | - | - | - | L248 |  | 7.146 | - | - | - | - | - | - | K296 |  | 2.66 | - | - | - |
| - | - | - | - | - | - | M250 |  | 3.089 | - | - | - | - | - | - | - | - | - | - | - | - |
| - | - | - | - | - | - | T252 |  | 5.94 | - | - | - | - | - | - | - | - | - | - | - | - |
| - | - | - | - | - | - | N289 |  | 7.061 | - | - | - | - | - | - | - | - | - | - | - | - |
| - | - | - | - | - | - | T291 |  | 5.16 | - | - | - | - | - | - | - | - | - | - | - | - |
| - | - | - | - | - | - | N293 |  | 7.435 | - | - | - | - | - | - | - | - | - | - | - | - |
| - | - | - | - | - | - | K296 |  | 7.244 | - | - | - | - | - | - | - | - | - | - | - | - |

**Table S3.** Binding affinity of p-RBD of B21R protein with CRD region of DC-SIGN.

| Complex | ΔG (kcal mol^-1^) | K_d_ (M) at 25℃ | ICs charged-charged | ICs charged-polar | ICs charged-apolar | ICs polar-polar | ICs polar-apolar | ICs apolar-apolar | NIS charged | NIS apolar | Intermolecular contacts |
| --- | --- | --- | --- | --- | --- | --- | --- | --- | --- | --- | --- |
| 1996 | -12.4 | 7.9e-10 | 10.0 | 27.0 | 22.0 | 12.0 | 21.0 | 7.0 | 20.99 | 33.15 | 99 |
| 2003 | -13.9 | 6.1e-11 | 11.0 | 25.0 | 30.0 | 11.0 | 24.0 | 9.0 | 20.27 | 35.07 | 110 |
| 2017 | -14.2 | 3.6e-11 | 17.0 | 25.0 | 29.0 | 13.0 | 24.0 | 11.0 | 20.72 | 33.34 | 119 |
| 2022 | -12.9 | 3.6e-10 | 10.0 | 22.0 | 27.0 | 16.0 | 24.0 | 8.0 | 19.23 | 34.07 | 107 |
| 2023 | -12.0 | 1.7e-09 | 9.0 | 22.0 | 22.0 | 7.0 | 15.0 | 14.0 | 20.56 | 33.33 | 89 |
| 2024 | -11.8 | 2.2e-09 | 8.0 | 21.0 | 15.0 | 13.0 | 23.0 | 7.0 | 20.22 | 33.52 | 87 |
| 2025 | -19.3 | 6.6e-15 | 14.0 | 28.0 | 44.0 | 17.0 | 44.0 | 33.0 | 20.34 | 33.05 | 180 |

**Table S4**. N-linked glycosylation sites in the B21R protein of MPVX strains from 1996 to 2025**.**

| Position | Sequence | Potential | Jury Agreement | N-Glyc Result |
| --- | --- | --- | --- | --- |
| Zaire-96-I-16-B21R | | | | |
| 69 | NWTD | 0.6552 | (9/9) | ++ |
| 94 | NYTI | 0.7724 | (9/9) | +++ |
| 124 | NYTI | 0.7724 | (9/9) | +++ |
| 227 | NVSV | 0.7312 | (9/9) | ++ |
| 241 | NDTS | 0.7181 | (9/9) | ++ |
| 276 | NVSI | 0.7261 | (9/9) | ++ |
| 289 | NVTS | 0.7751 | (9/9) | +++ |
| 340 | NSTC | 0.6710 | (9/9) | ++ |
| 355 | NITL | 0.7018 | (9/9) | ++ |
| 1809 | NSSP | 0.0994 | (9/9) | --- |
| 2003-MPXV-B21R | | | | |
| 69 | NWTD | 0.6553 | (9/9) | ++ |
| 94 | NYTI | 0.7546 | (9/9) | +++ |
| 124 | NYTI | 0.7724 | (9/9) | +++ |
| 227 | NVSV | 0.7312 | (9/9) | ++ |
| 241 | NDTS | 0.7181 | (9/9) | ++ |
| 276 | NVSI | 0.7262 | (9/9) | ++ |
| 289 | NVTS | 0.7751 | (9/9) | +++ |
| 340 | NSTC | 0.6707 | (9/9) | ++ |
| 355 | NITL | 0.7018 | (9/9) | ++ |
| 1810 | NSSP | 0.0994 | (9/9) | --- |
| 2017-MPXV-B21R | | | | |
| 69 | NWTD | 0.6553 | (9/9) | ++ |
| 94 | NYTI | 0.7547 | (9/9) | +++ |
| 124 | NYTI | 0.7724 | (9/9) | +++ |
| 227 | NVSV | 0.7312 | (9/9) | ++ |
| 241 | NDTS | 0.7181 | (9/9) | ++ |
| 276 | NVSI | 0.7262 | (9/9) | ++ |
| 289 | NVTS | 0.7751 | (9/9) | +++ |
| 340 | NSTC | 0.6707 | (9/9) | ++ |
| 355 | NITL | 0.7122 | (9/9) | ++ |
| 1810 | NSSP | 0.0994 | (9/9) | --- |
| 2022-MPXV-B21R | | | | |
| 69 | NWTD | 0.6554 | (9/9) | ++ |
| 94 | NYTI | 0.7547 | (9/9) | +++ |
| 124 | NYTI | 0.7724 | (9/9) | +++ |
| 227 | NVSV | 0.7312 | (9/9) | ++ |
| 241 | NDTS | 0.7181 | (9/9) | ++ |
| 276 | NVSI | 0.7262 | (9/9) | ++ |
| 289 | NVTS | 0.7751 | (9/9) | +++ |
| 340 | NSTC | 0.6707 | (9/9) | ++ |
| 355 | NITL | 0.7122 | (9/9) | ++ |
| 1810 | NSSP | 0.0994 | (9/9) | --- |
| 2023-MPXV-B21R | | | | |
| 69 | NWTD | 0.6554 | (9/9) | ++ |
| 94 | NYTI | 0.7547 | (9/9) | +++ |
| 124 | NYTI | 0.7724 | (9/9) | +++ |
| 227 | NVSV | 0.7312 | (9/9) | ++ |
| 241 | NDTS | 0.7181 | (9/9) | ++ |
| 276 | NVSI | 0.7262 | (9/9) | ++ |
| 289 | NVTS | 0.7751 | (9/9) | +++ |
| 340 | NSTC | 0.6707 | (9/9) | ++ |
| 355 | NITL | 0.7122 | (9/9) | ++ |
| 1810 | NSSP | 0.0994 | (9/9) | --- |
| 2024-MPXV-B21R | | | | |
| 69 | NWTD | 0.6554 | (9/9) | ++ |
| 94 | NYTI | 0.7547 | (9/9) | +++ |
| 124 | NYTI | 0.7724 | (9/9) | +++ |
| 227 | NVSV | 0.7312 | (9/9) | ++ |
| 241 | NDTS | 0.7181 | (9/9) | ++ |
| 276 | NVSI | 0.7262 | (9/9) | ++ |
| 289 | NVTS | 0.7751 | (9/9) | +++ |
| 340 | NSTC | 0.6707 | (9/9) | ++ |
| 355 | NITL | 0.7122 | (9/9) | ++ |
| 1810 | NSSP | 0.0994 | (9/9) | --- |
| 2025-MPXV-B21R | | | | |
| 69 | NWTD | 0.6553 | (9/9) | ++ |
| 94 | NYTI | 0.7547 | (9/9) | +++ |
| 124 | NYTI | 0.7724 | (9/9) | +++ |
| 227 | NVSV | 0.7312 | (9/9) | ++ |
| 241 | NDTS | 0.7181 | (9/9) | ++ |
| 276 | NVSI | 0.7262 | (9/9) | ++ |
| 289 | NVTS | 0.7751 | (9/9) | +++ |
| 340 | NSTC | 0.6707 | (9/9) | ++ |
| 355 | NITL | 0.7122 | (9/9) | ++ |
| 1810 | NSSP | 0.0994 | (9/9) | --- |

**Table S5.** O-linked glycosylation sites in the B21R protein of MPVX strains from 1996 to 2025.

| Site | Score | Site | Score | Site | Score | Site | Score | Site | Score | Site | Score | Site | Score |
| --- | --- | --- | --- | --- | --- | --- | --- | --- | --- | --- | --- | --- | --- |
| Zaire-96-I-16-B21R | | **2003-MPXV-B21R** | | **2017-MPXV-B21R** | | **2022-MPXV-B21R** | | **2023-MPXV-B21R** | | **2024-MPXV-B21R** | | **2025-MPXV-B21R** | |
| - | - | 47 | 0.5178 | - | - | - | - | - | - | - | - | - | - |
| 141 | 0.63005 | 141 | 0.6214 | 141 | 0.623 | 141 | 0.65 | 141 | 0.65 | 141 | 0.65 | 141 | 0.624257 |
| 143 | 0.84824 | 143 | 0.8624 | 143 | 0.8442 | 143 | 0.8499 | 143 | 0.8499 | 143 | 0.8499 | 143 | 0.844834 |
| 148 | 0.86014 | 148 | 0.8652 | 148 | 0.8452 | 148 | 0.8534 | 148 | 0.8534 | 148 | 0.8534 | 148 | 0.845854 |
| 149 | 0.89842 | 149 | 0.889 | 149 | 0.8906 | 149 | 0.9014 | 149 | 0.9014 | 149 | 0.9014 | 149 | 0.890961 |
| 153 | 0.82596 | 153 | 0.846 | 153 | 0.8112 | 153 | 0.8425 | 153 | 0.8425 | 153 | 0.8425 | 153 | 0.812105 |
| 154 | 0.88301 | 154 | 0.8953 | 154 | 0.8642 | 154 | 0.8885 | 154 | 0.8885 | 154 | 0.8885 | 154 | 0.864793 |
| 365 | 0.92538 | 365 | 0.9053 | 365 | 0.8932 | 365 | 0.8845 | 365 | 0.8845 | 365 | 0.8845 | 365 | 0.893508 |
| 367 | 0.76761 | 367 | 0.6917 | 367 | 0.7314 | 367 | 0.7285 | 367 | 0.7285 | 367 | 0.7285 | 367 | 0.731859 |
| 368 | 0.74807 | 368 | 0.6594 | 368 | 0.7099 | 368 | 0.6989 | 368 | 0.6989 | 368 | 0.6989 | 368 | 0.710259 |
| 369 | 0.92226 | 369 | 0.8881 | 369 | 0.9186 | 369 | 0.9165 | 369 | 0.9165 | 369 | 0.9165 | 369 | 0.918661 |
| 371 | 0.76552 | 371 | 0.5408 | 371 | 0.6776 | 371 | 0.6838 | 371 | 0.6838 | 371 | 0.6838 | 371 | 0.678009 |
| 384 | 0.62296 | 384 | 0.5 | - | - |  |  |  |  |  |  |  |  |
| 408 | 0.84694 | 408 | 0.8825 | 408 | 0.857 | 408 | 0.8525 | 408 | 0.8525 | 408 | 0.8525 | 408 | 0.857258 |
| 414 | 0.84731 | 414 | 0.9173 | 414 | 0.8665 | 414 | 0.872 | 414 | 0.872 | 414 | 0.872 | 414 | 0.866722 |
| 487 | 0.5098 | 487 | 0.6207 | 487 | 0.5174 | 487 | 0.5179 | 487 | 0.5179 | 487 | 0.5179 | 487 | 0.518174 |
| 494 | 0.58317 | 494 | 0.6342 | 494 | 0.5851 | 494 | 0.5905 | 494 | 0.5905 | 494 | 0.5905 | 494 | 0.585997 |
| 532 | 0.54596 | 532 | 0.6143 | 532 | 0.5649 | 532 | 0.5683 | 532 | 0.5683 | 532 | 0.5683 | 532 | 0.565821 |
| 539 | 0.53242 | 539 | 0.6272 | - | - | - | - | - | - | - | - | - | - |
| 546 | 0.67915 | 546 | 0.7963 | 546 | 0.6602 | 546 | 0.6687 | 546 | 0.6687 | 546 | 0.6687 | 546 | 0.660904 |
| 553 | 0.6375 | 553 | 0.6873 | 553 | 0.6511 | 553 | 0.6524 | 553 | 0.6524 | 553 | 0.6524 | 553 | 0.651906 |
| 557 | 0.67977 | 557 | 0.6824 | 557 | 0.672 | 557 | 0.677 | 557 | 0.677 | 557 | 0.677 | 557 | 0.676511 |
| 561 | 0.56415 | 561 | 0.6805 | 561 | 0.5668 | 561 | 0.5684 | 561 | 0.5684 | 561 | 0.5684 | 561 | 0.567628 |
| - | - | 563 | 0.5328 | - | - | - | - | - | - | - | - | - | - |
| 584 | 0.51468 | 584 | 0.6543 | 584 | 0.5156 | 584 | 0.5163 | 584 | 0.5163 | 584 | 0.5163 | 584 | 0.516894 |
| 588 | 0.69403 | 588 | 0.8184 | 588 | 0.6974 | 588 | 0.6922 | 588 | 0.6922 | 588 | 0.6922 | 588 | 0.698042 |
| 590 | 0.65751 | 590 | 0.8504 | 590 | 0.6697 | 590 | 0.67 | 590 | 0.67 | 590 | 0.67 | 590 | 0.670755 |
| - | - | 591 | 0.7525 | 591 | 0.5083 | 591 | 0.5099 | 591 | 0.5099 | 591 | 0.5099 | 591 | 0.509516 |
| 592 | 0.53481 | 592 | 0.8216 | 592 | 0.5391 | 592 | 0.5359 | 592 | 0.5359 | 592 | 0.5359 | 592 | 0.540271 |
| 593 | 0.74691 | 593 | 0.9038 | 593 | 0.7514 | 593 | 0.7487 | 593 | 0.7487 | 593 | 0.7487 | 593 | 0.752359 |
| - | - | 595 | 0.572 |  |  |  |  |  |  |  |  |  |  |
| 620 | 0.82452 | 620 | 0.939 | 620 | 0.8262 | 620 | 0.8239 | 620 | 0.8239 | 620 | 0.8239 | 620 | 0.826607 |
| 621 | 0.64049 | 621 | 0.8607 | 621 | 0.6358 | 621 | 0.6419 | 621 | 0.6419 | 621 | 0.6419 | 621 | 0.636366 |
| 622 | 0.83344 | 622 | 0.9263 | 622 | 0.8404 | 622 | 0.8401 | 622 | 0.8401 | 622 | 0.8401 | 622 | 0.840854 |
| 630 | 0.73063 | 630 | 0.9211 | 630 | 0.728 | 630 | 0.7276 | 630 | 0.7276 | 630 | 0.7276 | 630 | 0.728743 |
| - | - | 631 | 0.7981 | - | - | - | - | - | - | - | - | - | - |
| 635 | 0.5179 | 635 | 0.8303 | 635 | 0.5178 | 635 | 0.5208 | 635 | 0.5208 | 635 | 0.5208 | 635 | 0.518618 |
| 636 | 0.7667 | 636 | 0.9366 | 636 | 0.7675 | 636 | 0.774 | 636 | 0.774 | 636 | 0.774 | 636 | 0.768215 |
| 638 | 0.51028 | 638 | 0.8817 | 638 | 0.5137 | 638 | 0.5151 | 638 | 0.5151 | 638 | 0.5151 | 638 | 0.514679 |
| 640 | 0.55201 | 640 | 0.9143 | 640 | 0.5677 | 640 | 0.5697 | 640 | 0.5697 | 640 | 0.5697 | 640 | 0.568781 |
| - | - | 641 | 0.8018 | - | - | - | - | - | - | - | - | - | - |
| - | - | 647 | 0.7062 | - | - | - | - | - | - | - | - | - | - |
| - | - | 650 | 0.7866 | - | - | - | - | - | - | - | - | - | - |
| - | - | 653 | 0.5636 | - | - | - | - | - | - | - | - | - | - |
| - | - | 660 | 0.5201 | - | - | - | - | - | - | - | - | - | - |
| - | - | 662 | 0.6867 | - | - | - | - | - | - | - | - | - | - |
| 663 | 0.5 | 663 | 0.9044 | 663 | 0.5164 | 663 | 0.5257 | 663 | 0.5257 | 663 | 0.5257 | 663 | 0.517811 |
| 666 | 0.68931 | 666 | 0.9415 | 666 | 0.6924 | 666 | 0.6993 | 666 | 0.6993 | 666 | 0.6993 | 666 | 0.693384 |
| 669 | 0.58946 | 669 | 0.9043 | 669 | 0.5942 | 669 | 0.6004 | 669 | 0.6004 | 669 | 0.6004 | 669 | 0.595611 |
| 674 | 0.9688 | 674 | 0.9927 | 674 | 0.9691 | 674 | 0.9691 | 674 | 0.9691 | 674 | 0.9691 | 674 | 0.969218 |
| 682 | 0.94208 | 682 | 0.9751 | 682 | 0.9451 | 682 | 0.9472 | 682 | 0.9472 | 682 | 0.9472 | 682 | 0.945238 |
| 688 | 0.95158 | 688 | 0.9659 | 688 | 0.9527 | 688 | 0.9543 | 688 | 0.9543 | 688 | 0.9543 | 688 | 0.952834 |
| 704 | 0.83107 | - | - | 704 | 0.8552 | 704 | 0.842 | 704 | 0.842 | 704 | 0.842 | 704 | 0.85547 |
| 713 | 0.97689 | - | - | - | - | - | - | - | - | - | - | - | - |
| - | - | - | - | - | - | 722 | 0.9283 | 722 | 0.9283 | 722 | 0.9283 | - | - |
| 723 | 0.94386 | - | - | - | - | - | - | - | - | - | - | - | - |
| 745 | 0.83021 | - | - | - | - | - | - | - | - | - | - | - | - |
| - | - | 746 | 0.8929 | 746 | 0.8123 | 746 | 0.8081 | 746 | 0.8081 | 746 | 0.8081 | 746 | 0.813045 |
| 749 | 0.70346 | - | - | - | - | - | - | - | - | - | - | - | - |
| - | - | 750 | 0.8835 | 750 | 0.6924 | 750 | 0.6874 | 750 | 0.6874 | 750 | 0.6874 | 750 | 0.693007 |
| - | - | 758 | 0.7566 | - | - | - | - | - | - | - | - |  |  |
| - | - | 759 | 0.5386 | - | - | - | - | - | - | - | - |  |  |
| - | - | 776 | 0.5615 | - | - | - | - | - | - | - | - |  |  |
| 806 | 0.739 | - | - | - | - | - | - | - | - | - | - |  |  |
| - | - | 807 | 0.8019 | 807 | 0.7263 | 807 | 0.7278 | 807 | 0.7278 | 807 | 0.7278 | 807 | 0.726885 |
| 810 | 0.52597 | - | - | - | - | - | - | - | - | - | - |  |  |
| 811 | 0.70771 | 811 | 0.6348 | 811 | 0.5099 | 811 | 0.5117 | 811 | 0.5117 | 811 | 0.5117 | 811 | 0.510785 |
| - | - | 812 | 0.7746 | 812 | 0.6949 | 812 | 0.6964 | 812 | 0.6964 | 812 | 0.6964 | 812 | 0.695784 |
| 842 | 0.60763 | - | - | - | - | - | - | - | - | - | - |  |  |
| - | - | 843 | 0.566 | 843 | 0.6061 | 843 | 0.6119 | 843 | 0.6119 | 843 | 0.6119 | 843 | 0.607405 |
| 844 | 0.52959 | - | - | - | - | - | - | - | - | - | - |  |  |
| 845 | 0.56869 | 845 | 0.5262 | 845 | 0.5295 | 845 | 0.5364 | 845 | 0.5364 | 845 | 0.5364 | 845 | 0.53072 |
| 846 | 0.746 | 846 | 0.5434 | 846 | 0.5675 | 846 | 0.5704 | 846 | 0.5704 | 846 | 0.5704 | 846 | 0.568749 |
|  |  | 847 | 0.6912 | 847 | 0.7436 | 847 | 0.7478 | 847 | 0.7478 | 847 | 0.7478 | 847 | 0.744707 |
| 848 | 0.55785 | - | - | - | - | - | - | - | - | - | - |  |  |
| - | - | 849 | 0.5379 | 849 | 0.5531 | 849 | 0.5643 | 849 | 0.5643 | 849 | 0.5643 | 849 | 0.55439 |
| 1069 | 0.53068 | - | - | - | - | - | - | - | - | - | - |  |  |
| - | - | - | - | 1070 | 0.5378 | 1070 | 0.5389 | 1070 | 0.5389 | 1070 | 0.5389 | 1070 | 0.538311 |
| 1075 | 0.62032 | - | - | - | - | - | - | - | - | - | - |  |  |
| 1076 | 0.62757 | 1076 | 0.5787 | 1076 | 0.6136 | 1076 | 0.6151 | 1076 | 0.6151 | 1076 | 0.6151 | 1076 | 0.614234 |
| - | - | 1077 | 0.5917 | 1077 | 0.6296 | 1077 | 0.631 | 1077 | 0.631 | 1077 | 0.631 | 1077 | 0.630212 |
| 1305 | 0.74808 | - | - | - | - | - | - | - | - | - | - |  |  |
| - | - | 1306 | 0.6683 | 1306 | 0.7497 | 1306 | 0.7508 | 1306 | 0.7508 | 1306 | 0.7508 | 1306 | 0.750288 |
| 1314 | 0.62047 | - | - | - | - | - | - | - | - | - | - | - | - |
| - | - | 1315 | 0.5833 | 1315 | 0.642 | 1315 | 0.6432 | 1315 | 0.6432 | 1315 | 0.6432 | 1315 | 0.642352 |
| 1513 | 0.68167 | - | - | - | - | - | - | - | - | - | - | - | - |
| - | - | 1514 | 0.6573 | 1514 | 0.6926 | 1514 | 0.6932 | 1514 | 0.6932 | 1514 | 0.6932 | 1514 | 0.693058 |

Table S6. CTL epitopes of B21R protein of MPVX strains from 1996 to 2025.

| POSITION | PEPTIDE | AFF | AFF_RESCALE | CLEAVAGE | TAP | COMB |
| --- | --- | --- | --- | --- | --- | --- |
| Zaire-96-I-16-B21R | | | | | | |
| 12 | LTVTCSWCY | 0.6345 | 2.6940 | 0.9382 | 2.9330 | 2.9814 <-E |
| 44 | SVASLPYKY | 0.2833 | 1.2029 | 0.7983 | 2.9940 | 1.4723 <-E |
| 70 | WTDIAEGVR | 0.1693 | 0.7187 | 0.0432 | 1.4010 | 0.7952 <-E |
| 83 | KICDINGTY | 0.3071 | 1.3038 | 0.9093 | 3.1780 | 1.5991 <-E |
| 110 | PTVTPITTY | 0.2319 | 0.9844 | 0.9723 | 2.5490 | 1.2577 <-E |
| 115 | ITTYEPSIY | 0.4981 | 2.1148 | 0.6249 | 2.7450 | 2.3458 <-E |
| 117 | TYEPSIYNY | 0.1394 | 0.5919 | 0.9741 | 3.0340 | 0.8897 <-E |
| 121 | SIYNYTIDY | 0.2145 | 0.9106 | 0.9727 | 3.1480 | 1.2139 <-E |
| 182 | ETELTNYLI | 0.2056 | 0.8731 | 0.6316 | 0.3060 | 0.9831 <-E |
| 206 | ETSDNNTLH | 0.2687 | 1.1410 | 0.1596 | 0.9040 | 1.1197 <-E |
| 207 | TSDNNTLHG | 0.2184 | 0.9274 | 0.1116 | -1.7190 | 0.8582 <-E |
| 242 | DTSEEILLM | 0.2115 | 0.8980 | 0.8967 | 0.1040 | 1.0377 <-E |
| 243 | TSEEILLML | 0.2014 | 0.8550 | 0.9652 | 0.6910 | 1.0343 <-E |
| 256 | SSDTFISST | 0.2687 | 1.1406 | 0.3474 | -0.7340 | 1.1561 <-E |
| 266 | ITECLKTLI | 0.1650 | 0.7005 | 0.2384 | 0.5330 | 0.7629 <-E |
| 322 | KDDENNTVY | 0.1514 | 0.6429 | 0.9210 | 2.5950 | 0.9108 <-E |
| 325 | ENNTVYNYY | 0.1202 | 0.5103 | 0.8782 | 2.3350 | 0.7588 <-E |
| 385 | TSKELDCLY | 0.4246 | 1.8030 | 0.8142 | 2.9960 | 2.0749 <-E |
| 388 | ELDCLYESY | 0.5389 | 2.2882 | 0.9343 | 2.6650 | 2.5616 <-E |
| 413 | RSDDKKEYM | 0.1490 | 0.6327 | 0.6499 | 0.5450 | 0.7575 <-E |
| 420 | YMDMKLFDH | 0.1853 | 0.7867 | 0.1055 | -0.7770 | 0.7637 <-E |
| 484 | ITATEADLY | 0.7400 | 3.1419 | 0.5413 | 2.7980 | 3.3630 <-E |
| 510 | FTEALASTI | 0.2634 | 1.1182 | 0.1995 | 0.1290 | 1.1546 <-E |
| 522 | LSNVREVTY | 0.3623 | 1.5382 | 0.9725 | 2.9760 | 1.8329 <-E |
| 569 | DIQTVVKEY | 0.1160 | 0.4924 | 0.9665 | 2.5380 | 0.7643 <-E |
| 573 | VVKEYNERY | 0.1967 | 0.8353 | 0.9655 | 3.1330 | 1.1368 <-E |
| 599 | DIDTVVREY | 0.4725 | 2.0063 | 0.9535 | 2.3940 | 2.2690 <-E |
| 621 | SSPKPDPLY | 0.2237 | 0.9499 | 0.9712 | 3.0570 | 1.2485 <-E |
| 644 | DIVTKQSDY | 0.1120 | 0.4754 | 0.9624 | 2.8620 | 0.7629 <-E |
| 766 | QSKPNDDTY | 0.2887 | 1.2260 | 0.9719 | 2.9850 | 1.5210 <-E |
| 812 | TMSKISTKF | 0.1118 | 0.4747 | 0.9745 | 2.6850 | 0.7551 <-E |
| 898 | IIDTIKDIY | 0.5021 | 2.1320 | 0.7047 | 2.8700 | 2.3812 <-E |
| 930 | MSDNNKMGV | 0.3502 | 1.4867 | 0.9273 | 0.2740 | 1.6395 <-E |
| 962 | QNTDAMALY | 0.1404 | 0.5962 | 0.9208 | 2.8500 | 0.8769 <-E |
| 963 | NTDAMALYF | 0.7012 | 2.9773 | 0.2818 | 2.3250 | 3.1359 <-E |
| 971 | FLDVIDSEI | 0.1719 | 0.7298 | 0.5389 | 0.2310 | 0.8221 <-E |
| 973 | DVIDSEILY | 0.1294 | 0.5494 | 0.9420 | 2.8640 | 0.8339 <-E |
| 974 | VIDSEILYL | 0.1505 | 0.6388 | 0.8876 | 0.8300 | 0.8135 <-E |
| 981 | YLNTSNLVL | 0.1464 | 0.6215 | 0.9097 | 0.9080 | 0.8034 <-E |
| 983 | NTSNLVLEY | 0.7968 | 3.3830 | 0.9685 | 2.9840 | 3.6774 <-E |
| 1004 | SVDVDITAY | 0.6239 | 2.6490 | 0.9691 | 3.0720 | 2.9479 <-E |
| 1008 | DITAYTILY | 0.3332 | 1.4148 | 0.9740 | 2.7480 | 1.6983 <-E |
| 1017 | DTADNIKKY | 0.3513 | 1.4917 | 0.9417 | 2.5520 | 1.7606 <-E |
| 1059 | ISDMQLLKM | 0.2133 | 0.9057 | 0.9513 | 0.0470 | 1.0507 <-E |
| 1093 | YLLAGGCPY | 0.1462 | 0.6206 | 0.9726 | 2.8370 | 0.9084 <-E |
| 1110 | HTTCSILLR | 0.1404 | 0.5959 | 0.6701 | 1.3830 | 0.7656 <-E |
| 1191 | CSSRTRKIY | 0.2490 | 1.0571 | 0.1648 | 2.9740 | 1.2305 <-E |
| 1237 | IIELPVGDY | 0.1955 | 0.8299 | 0.9433 | 2.9790 | 1.1203 <-E |
| 1251 | YSATKPSRI | 0.1634 | 0.6938 | 0.2680 | 0.6200 | 0.7650 <-E |
| 1259 | IAVFCTHNY | 0.1058 | 0.4492 | 0.9573 | 3.2100 | 0.7533 <-E |
| 1272 | KSDIIVLMF | 0.5623 | 2.3873 | 0.9337 | 2.4010 | 2.6474 <-E |
| 1335 | DSVEADIHY | 0.2578 | 1.0945 | 0.7908 | 2.5600 | 1.3411 <-E |
| 1379 | MVDNLGNGY | 0.6877 | 2.9200 | 0.8658 | 2.8290 | 3.1914 <-E |
| 1436 | YLQSTSQDY | 0.3379 | 1.4345 | 0.9011 | 2.9390 | 1.7167 <-E |
| 1481 | SIPRNISTY | 0.1287 | 0.5464 | 0.9758 | 2.9860 | 0.8420 <-E |
| 1524 | RACFHHWNY | 0.3270 | 1.3883 | 0.8099 | 3.0850 | 1.6641 <-E |
| 1531 | NYYTLSLDY | 0.1121 | 0.4761 | 0.9718 | 3.3570 | 0.7897 <-E |
| 1532 | YYTLSLDYY | 0.1099 | 0.4666 | 0.9469 | 3.2740 | 0.7724 <-E |
| 1535 | LSLDYYCSY | 0.4286 | 1.8198 | 0.8891 | 2.9970 | 2.1030 <-E |
| 1557 | CKSYIHIEY | 0.1989 | 0.8446 | 0.9262 | 2.7840 | 1.1227 <-E |
| 1584 | FIHDNSNEY | 0.3652 | 1.5507 | 0.9708 | 2.8210 | 1.8374 <-E |
| 1597 | ISNKLNDLY | 0.6218 | 2.6403 | 0.4412 | 2.7270 | 2.8428 <-E |
| 1600 | KLNDLYNEY | 0.3068 | 1.3026 | 0.9555 | 3.0980 | 1.6008 <-E |
| 1713 | YLCGTLDDY | 0.2159 | 0.9168 | 0.3733 | 3.0740 | 1.1265 <-E |
| 1717 | TLDDYIYTV | 0.2191 | 0.9302 | 0.9744 | 0.1070 | 1.0817 <-E |
| 1724 | TVVEYNKSY | 0.1340 | 0.5689 | 0.9569 | 3.0680 | 0.8659 <-E |
| 1734 | LVNDTYMSY | 0.3559 | 1.5111 | 0.9778 | 3.0200 | 1.8088 <-E |
| 1807 | YLNSSPDIY | 0.4086 | 1.7347 | 0.4269 | 2.9710 | 1.9472 <-E |
| 1811 | SPDIYHIIY | 0.2862 | 1.2153 | 0.9699 | 2.5510 | 1.4883 <-E |
| 1871 | SMEIIDNRY | 0.5615 | 2.3839 | 0.9756 | 3.0020 | 2.6803 <-E |
| 2003-MPXV-B21R | | | | | | |
| 12 | LTVTCSWCY | 0.6345 | 2.6940 | 0.9382 | 2.9330 | 2.9814 <-E |
| 44 | SVASLPYKY | 0.2833 | 1.2029 | 0.7983 | 2.9940 | 1.4723 <-E |
| 70 | WTDIAEGVR | 0.1693 | 0.7187 | 0.0432 | 1.4010 | 0.7952 <-E |
| 83 | KICDINGTY | 0.3071 | 1.3038 | 0.8688 | 3.1780 | 1.5930 <-E |
| 110 | PTVTPITTY | 0.2319 | 0.9844 | 0.9723 | 2.5490 | 1.2577 <-E |
| 117 | TYEPSIYNY | 0.1394 | 0.5919 | 0.9741 | 3.0340 | 0.8897 <-E |
| 121 | SIYNYTIDY | 0.2145 | 0.9106 | 0.9727 | 3.1480 | 1.2139 <-E |
| 115 | ITTYEPSIY | 0.4981 | 2.1148 | 0.6249 | 2.7450 | 2.3458 <-E |
| 182 | ETELTNYLI | 0.2056 | 0.8731 | 0.6316 | 0.3060 | 0.9831 <-E |
| 207 | TSDNNTLHG | 0.2184 | 0.9274 | 0.1116 | -1.7190 | 0.8582 <-E |
| 206 | ETSDNNTLH | 0.2687 | 1.1410 | 0.1596 | -0.9040 | 1.1197 <-E |
| 242 | DTSEEILLM | 0.2115 | 0.8980 | 0.8967 | 0.1040 | 1.0377 <-E |
| 243 | TSEEILLML | 0.2014 | 0.8550 | 0.9652 | 0.6910 | 1.0343 <-E |
| 256 | SSDTFISST | 0.2687 | 1.1406 | 0.3474 | -0.7340 | 1.1561 <-E |
| 266 | ITECLKTLI | 0.1650 | 0.7005 | 0.2384 | 0.5330 | 0.7629 <-E |
| 322 | KDDENNTVY | 0.1514 | 0.6429 | 0.9539 | 2.5950 | 0.9157 <-E |
| 385 | TSKELDCLY | 0.4246 | 1.8030 | 0.8142 | 2.9960 | 2.0749 <-E |
| 388 | ELDCLYESY | 0.5389 | 2.2882 | 0.9343 | 2.6650 | 2.5616 <-E |
| 413 | RSDDKKEYM | 0.1490 | 0.6327 | 0.6499 | 0.5450 | 0.7575 <-E |
| 420 | YMDMKLFDH | 0.1853 | 0.7867 | 0.1055 | -0.7770 | 0.7637 <-E |
| 484 | ITATEADLY | 0.7400 | 3.1419 | 0.5413 | 2.7980 | 3.3630 <-E |
| 510 | FTEALVSTI | 0.2466 | 1.0469 | 0.3170 | 0.1290 | 1.1009 <-E |
| 522 | LSNVREVTY | 0.3623 | 1.5382 | 0.9674 | 2.9760 | 1.8321 <-E |
| 569 | DIQTVVKEY | 0.1160 | 0.4924 | 0.9665 | 2.5380 | 0.7643 <-E |
| 573 | VVKEYNERY | 0.1967 | 0.8353 | 0.9655 | 3.1330 | 1.1368 <-E |
| 599 | DIDTVVREY | 0.4725 | 2.0063 | 0.9535 | 2.3940 | 2.2690 <-E |
| 621 | SSPKPDPLY | 0.2237 | 0.9499 | 0.9712 | 3.0570 | 1.2485 <-E |
| 644 | DIVTKQSDY | 0.1120 | 0.4754 | 0.9624 | 2.8620 | 0.7629 <-E |
| 767 | QSKPNDDTY | 0.2887 | 1.2260 | 0.9719 | 2.9850 | 1.5210 <-E |
| 813 | TMSKISTKF | 0.1118 | 0.4747 | 0.9745 | 2.6850 | 0.7551 <-E |
| 899 | IIDTIKDIY | 0.5021 | 2.1320 | 0.7047 | 2.8700 | 2.3812 <-E |
| 931 | MSDNNKMGV | 0.3502 | 1.4867 | 0.9273 | 0.2740 | 1.6395 <-E |
| 963 | QNTDAMALY | 0.1404 | 0.5962 | 0.9208 | 2.8500 | 0.8769 <-E |
| 964 | NTDAMALYF | 0.7012 | 2.9773 | 0.2818 | 2.3250 | 3.1359 <-E |
| 972 | FLDVIDSEI | 0.1719 | 0.7298 | 0.5389 | 0.2310 | 0.8221 <-E |
| 974 | DVIDSEILY | 0.1294 | 0.5494 | 0.9420 | 2.8640 | 0.8339 <-E |
| 975 | VIDSEILYL | 0.1505 | 0.6388 | 0.8876 | 0.8300 | 0.8135 <-E |
| 982 | YLNTSNLVL | 0.1464 | 0.6215 | 0.9097 | 0.9080 | 0.8034 <-E |
| 984 | NTSNLVLEY | 0.7968 | 3.3830 | 0.9685 | 2.9840 | 3.6774 <-E |
| 1005 | SVDVDITAY | 0.6239 | 2.6490 | 0.9691 | 3.0720 | 2.9479 <-E |
| 1009 | DITAYTILY | 0.3332 | 1.4148 | 0.9740 | 2.7480 | 1.6983 <-E |
| 1018 | DTADNIKKY | 0.3513 | 1.4917 | 0.9417 | 2.5520 | 1.7606 <-E |
| 1060 | ISDMQLLKM | 0.2133 | 0.9057 | 0.9513 | 0.0470 | 1.0507 <-E |
| 1094 | YLLAGGCPY | 0.1462 | 0.6206 | 0.9726 | 2.8370 | 0.9084 <-E |
| 1111 | HTTCSILLR | 0.1404 | 0.5959 | 0.6701 | 1.3830 | 0.7656 <-E |
| 1192 | CSSRTRKIY | 0.2490 | 1.0571 | 0.1648 | 2.9740 | 1.2305 <-E |
| 1238 | IIELPVGDY | 0.1955 | 0.8299 | 0.9433 | 2.9790 | 1.1203 <-E |
| 1252 | YSATKPSRI | 0.1634 | 0.6938 | 0.2680 | 0.6200 | 0.7650 <-E |
| 1260 | IAVFCTHNY | 0.1058 | 0.4492 | 0.9573 | 3.2100 | 0.7533 <-E |
| 1273 | KSDIIVLMF | 0.5623 | 2.3873 | 0.9337 | 2.4010 | 2.6474 <-E |
| 1336 | DSVETDIHY | 0.2547 | 1.0813 | 0.8074 | 2.5600 | 1.3304 <-E |
| 1380 | MVDNLGNGY | 0.6877 | 2.9200 | 0.8658 | 2.8290 | 3.1914 <-E |
| 1437 | YLQSTSQDY | 0.3379 | 1.4345 | 0.9011 | 2.9390 | 1.7167 <-E |
| 1482 | SIPRNISTY | 0.1287 | 0.5464 | 0.9758 | 2.9860 | 0.8420 <-E |
| 1525 | RACFHHWNY | 0.3270 | 1.3883 | 0.8099 | 3.0850 | 1.6641 <-E |
| 1532 | NYYTLSLDY | 0.1121 | 0.4761 | 0.9718 | 3.3570 | 0.7897 <-E |
| 1533 | YYTLSLDYY | 0.1099 | 0.4666 | 0.9469 | 3.2740 | 0.7724 <-E |
| 1536 | LSLDYYCSY | 0.4286 | 1.8198 | 0.8891 | 2.9970 | 2.1030 <-E |
| 1558 | CKSYIHIEY | 0.1989 | 0.8446 | 0.9262 | 2.7840 | 1.1227 <-E |
| 1585 | FIHDNSNEY | 0.3652 | 1.5507 | 0.9708 | 2.8210 | 1.8374 <-E |
| 1598 | ISNKLNDLY | 0.6218 | 2.6403 | 0.4412 | 2.7270 | 2.8428 <-E |
| 1601 | KLNDLYNEY | 0.3068 | 1.3026 | 0.9555 | 3.0980 | 1.6008 <-E |
| 1697 | IIDVKNNLV | 0.1916 | 0.8135 | 0.5823 | 0.2830 | 0.9150 <-E |
| 1714 | YLCDTLDDY | 0.3048 | 1.2941 | 0.6966 | 3.0740 | 1.5523 <-E |
| 1725 | TSVEYNKSY | 0.2700 | 1.1465 | 0.9412 | 2.9700 | 1.4362 <-E |
| 1735 | LVNDTYMSY | 0.3559 | 1.5111 | 0.9778 | 3.0200 | 1.8088 <-E |
| 1808 | YLNSSPDIY | 0.4086 | 1.7347 | 0.4269 | 2.9710 | 1.9472 <-E |
| 1812 | SPDIYHIIY | 0.2862 | 1.2153 | 0.9699 | 2.5510 | 1.4883 <-E |
| 1872 | SMEIIDNRY | 0.5615 | 2.3839 | 0.9756 | 3.0020 | 2.6803 <-E |
| 2017-MPXV-B21R | | | | | | |
| 12 | LTVTCSWCY | 0.6345 | 2.6940 | 0.9382 | 2.9330 | 2.9814 <-E |
| 44 | SVASLPYKY | 0.2833 | 1.2029 | 0.7983 | 2.9940 | 1.4723 <-E |
| 70 | WTDIAEGVR | 0.1693 | 0.7187 | 0.0432 | 1.4010 | 0.7952 <-E |
| 83 | KICDINGTY | 0.3071 | 1.3038 | 0.8688 | 3.1780 | 1.5930 <-E |
| 110 | PTVTPITTY | 0.2319 | 0.9844 | 0.9723 | 2.5490 | 1.2577 <-E |
| 115 | ITTYEPSIY | 0.4981 | 2.1148 | 0.6249 | 2.7450 | 2.3458 <-E |
| 117 | TYEPSIYNY | 0.1394 | 0.5919 | 0.9741 | 3.0340 | 0.8897 <-E |
| 121 | SIYNYTIDY | 0.2145 | 0.9106 | 0.9727 | 3.1480 | 1.2139 <-E |
| 182 | ETELTNYLI | 0.2056 | 0.8731 | 0.6316 | 0.3060 | 0.9831 <-E |
| 206 | ETSDNNTLH | 0.2687 | 1.1410 | 0.1596 | -0.9040 | 1.1197 <-E |
| 207 | TSDNNTLHG | 0.2184 | 0.9274 | 0.1116 | -1.7190 | 0.8582 <-E |
| 242 | DTSEEILLM | 0.2115 | 0.8980 | 0.8967 | 0.1040 | 1.0377 <-E |
| 243 | TSEEILLML | 0.2014 | 0.8550 | 0.9652 | 0.6910 | 1.0343 <-E |
| 256 | SSDTFISST | 0.2687 | 1.1406 | 0.3474 | -0.7340 | 1.1561 <-E |
| 266 | ITECLKTLI | 0.1650 | 0.7005 | 0.2384 | 0.5330 | 0.7629 <-E |
| 322 | KDDENNTVY | 0.1514 | 0.6429 | 0.9539 | 2.5950 | 0.9157 <-E |
| 358 | LTNIIHNSV | 0.1606 | 0.6820 | 0.9502 | 0.2130 | 0.8352 <-E |
| 385 | TSKELDCLY | 0.4246 | 1.8030 | 0.8142 | 2.9960 | 2.0749 <-E |
| 388 | ELDCLYESY | 0.5389 | 2.2882 | 0.9343 | 2.6650 | 2.5616 <-E |
| 413 | RSDDKKEYM | 0.1490 | 0.6327 | 0.6499 | 0.5450 | 0.7575 <-E |
| 420 | YMDMKLFDH | 0.1853 | 0.7867 | 0.1055 | -0.7770 | 0.7637 <-E |
| 484 | ITATEADLY | 0.7400 | 3.1419 | 0.5413 | 2.7980 | 3.3630 <-E |
| 510 | FTEALVSTI | 0.2466 | 1.0469 | 0.3170 | 0.1290 | 1.1009 <-E |
| 522 | LSNVREVTY | 0.3623 | 1.5382 | 0.9674 | 2.9760 | 1.8321 <-E |
| 569 | DIQTVVKEY | 0.1160 | 0.4924 | 0.9665 | 2.5380 | 0.7643 <-E |
| 573 | VVKEYNERY | 0.1967 | 0.8353 | 0.9655 | 3.1330 | 1.1368 <-E |
| 599 | DIDTVVREY | 0.4725 | 2.0063 | 0.9535 | 2.3940 | 2.2690 <-E |
| 621 | SSPKPDPLY | 0.2237 | 0.9499 | 0.9712 | 3.0570 | 1.2485 <-E |
| 644 | DIVTKQSDY | 0.1120 | 0.4754 | 0.9624 | 2.8620 | 0.7629 <-E |
| 767 | QSKPNDDTY | 0.2887 | 1.2260 | 0.9706 | 2.9850 | 1.5208 <-E |
| 813 | TMSKISTKF | 0.1118 | 0.4747 | 0.9745 | 2.6850 | 0.7551 <-E |
| 899 | IIDTIKDIY | 0.5021 | 2.1320 | 0.7047 | 2.8700 | 2.3812 <-E |
| 931 | MSDNNKMGV | 0.3502 | 1.4867 | 0.9273 | 0.2740 | 1.6395 <-E |
| 963 | QNTDAMALY | 0.1404 | 0.5962 | 0.9208 | 2.8500 | 0.8769 <-E |
| 964 | NTDAMALYF | 0.7012 | 2.9773 | 0.2818 | 2.3250 | 3.1359 <-E |
| 972 | FLDVIDSEI | 0.1719 | 0.7298 | 0.5389 | 0.2310 | 0.8221 <-E |
| 974 | DVIDSEILY | 0.1294 | 0.5494 | 0.9420 | 2.8640 | 0.8339 <-E |
| 975 | VIDSEILYL | 0.1505 | 0.6388 | 0.8876 | 0.8300 | 0.8135 <-E |
| 982 | YLNTSNLVL | 0.1464 | 0.6215 | 0.9097 | 0.9080 | 0.8034 <-E |
| 984 | NTSNLVLEY | 0.7968 | 3.3830 | 0.9685 | 2.9840 | 3.6774 <-E |
| 1005 | SVDVDITAY | 0.6239 | 2.6490 | 0.9691 | 3.0720 | 2.9479 <-E |
| 1009 | DITAYTILY | 0.3332 | 1.4148 | 0.9740 | 2.7480 | 1.6983 <-E |
| 1018 | DTADNIKKY | 0.3513 | 1.4917 | 0.9417 | 2.5520 | 1.7606 <-E |
| 1060 | ISDMQLLKM | 0.2133 | 0.9057 | 0.9513 | 0.0470 | 1.0507 <-E |
| 1094 | YLLAGGCPY | 0.1462 | 0.6206 | 0.9726 | 2.8370 | 0.9084 <-E |
| 1111 | HTTCSILLR | 0.1404 | 0.5959 | 0.6701 | 1.3830 | 0.7656 <-E |
| 1192 | CSSRTRKIY | 0.2490 | 1.0571 | 0.1648 | 2.9740 | 1.2305 <-E |
| 1238 | IIELPVGDY | 0.1955 | 0.8299 | 0.9433 | 2.9790 | 1.1203 <-E |
| 1252 | YSATKPSRI | 0.1634 | 0.6938 | 0.2680 | 0.6200 | 0.7650 <-E |
| 1260 | IAVFCTHNY | 0.1058 | 0.4492 | 0.9573 | 3.2100 | 0.7533 <-E |
| 1273 | KSDIIVLMF | 0.5623 | 2.3873 | 0.9337 | 2.4010 | 2.6474 <-E |
| 1336 | DSVETDIHY | 0.2547 | 1.0813 | 0.8074 | 2.5600 | 1.3304 <-E |
| 1380 | MVDNLGNGY | 0.6877 | 2.9200 | 0.8658 | 2.8290 | 3.1914 <-E |
| 1437 | YLQSTSQDY | 0.3379 | 1.4345 | 0.9011 | 2.9390 | 1.7167 <-E |
| 1482 | SIPRNISTY | 0.1287 | 0.5464 | 0.9758 | 2.9860 | 0.8420 <-E |
| 1525 | RACFHHWNY | 0.3270 | 1.3883 | 0.8099 | 3.0850 | 1.6641 <-E |
| 1532 | NYYTLSLDY | 0.1121 | 0.4761 | 0.9718 | 3.3570 | 0.7897 <-E |
| 1533 | YYTLSLDYY | 0.1099 | 0.4666 | 0.9469 | 3.2740 | 0.7724 <-E |
| 1536 | LSLDYYCSY | 0.4286 | 1.8198 | 0.8891 | 2.9970 | 2.1030 <-E |
| 1558 | CKSYIHIEY | 0.1989 | 0.8446 | 0.9262 | 2.7840 | 1.1227 <-E |
| 1585 | FIHDNSNEY | 0.3652 | 1.5507 | 0.9708 | 2.8210 | 1.8374 <-E |
| 1598 | ISNKLNDLY | 0.6218 | 2.6403 | 0.4412 | 2.7270 | 2.8428 <-E |
| 1601 | KLNDLYNEY | 0.3068 | 1.3026 | 0.9555 | 3.0980 | 1.6008 <-E |
| 1697 | IIDVKNNLV | 0.1916 | 0.8135 | 0.5823 | 0.2830 | 0.9150 <-E |
| 1714 | YLCDTLDDY | 0.3048 | 1.2941 | 0.6966 | 3.0740 | 1.5523 <-E |
| 1725 | TSVEYNKSY | 0.2700 | 1.1465 | 0.9412 | 2.9700 | 1.4362 <-E |
| 1735 | LVNDTYMSY | 0.3559 | 1.5111 | 0.9778 | 3.0200 | 1.8088 <-E |
| 1808 | YLNSSPDIY | 0.4086 | 1.7347 | 0.4269 | 2.9710 | 1.9472 <-E |
| 1812 | SPDIYHIIY | 0.2862 | 1.2153 | 0.9699 | 2.5510 | 1.4883 <-E |
| 1872 | SMEIIDNRY | 0.5615 | 2.3839 | 0.9756 | 3.0020 | 2.6803 <-E |
| 2022-MPXV-B21R | | | | | | |
| 12 | LTVTCSWCY | 0.6345 | 2.6940 | 0.9382 | 2.9330 | 2.9814 <-E |
| 44 | SVASLPYKY | 0.2833 | 1.2029 | 0.7983 | 2.9940 | 1.4723 <-E |
| 70 | WTDIAEGVR | 0.1693 | 0.7187 | 0.0432 | 1.4010 | 0.7952 <-E |
| 83 | KICDINGTY | 0.3071 | 1.3038 | 0.8688 | 3.1780 | 1.5930 <-E |
| 110 | PTVTPITTY | 0.2319 | 0.9844 | 0.9723 | 2.5490 | 1.2577 <-E |
| 115 | ITTYEPSIY | 0.4981 | 2.1148 | 0.6249 | 2.7450 | 2.3458 <-E |
| 117 | TYEPSIYNY | 0.1394 | 0.5919 | 0.9741 | 3.0340 | 0.8897 <-E |
| 121 | SIYNYTIDY | 0.2145 | 0.9106 | 0.9727 | 3.1480 | 1.2139 <-E |
| 182 | ETELTNYLI | 0.2056 | 0.8731 | 0.6316 | 0.3060 | 0.9831 <-E |
| 206 | ETSNNNTLH | 0.2680 | 1.1380 | 0.1142 | -0.9040 | 1.1100 <-E |
| 242 | DTSEEILLM | 0.2115 | 0.8980 | 0.8967 | 0.1040 | 1.0377 <-E |
| 243 | TSEEILLML | 0.2014 | 0.8550 | 0.9652 | 0.6910 | 1.0343 <-E |
| 256 | SSDTFISST | 0.2687 | 1.1406 | 0.3474 | -0.7340 | 1.1561 <-E |
| 266 | ITECLKTLI | 0.1650 | 0.7005 | 0.2384 | 0.5330 | 0.7629 <-E |
| 322 | KDDENNTVY | 0.1514 | 0.6429 | 0.9539 | 2.5950 | 0.9157 <-E |
| 358 | LTNIIHNSV | 0.1606 | 0.6820 | 0.9502 | 0.2130 | 0.8352 <-E |
| 385 | TSKELDCLY | 0.4246 | 1.8030 | 0.8142 | 2.9960 | 2.0749 <-E |
| 388 | ELDCLYESY | 0.5389 | 2.2882 | 0.9343 | 2.6650 | 2.5616 <-E |
| 413 | RSDDKKEYM | 0.1490 | 0.6327 | 0.6499 | 0.5450 | 0.7575 <-E |
| 420 | YMDMKLFDH | 0.1853 | 0.7867 | 0.1055 | -0.7770 | 0.7637 <-E |
| 484 | ITATEADLY | 0.7400 | 3.1419 | 0.5413 | 2.7980 | 3.3630 <-E |
| 510 | FTEALVSTI | 0.2466 | 1.0469 | 0.3170 | 0.1290 | 1.1009 <-E |
| 522 | LSNVREVTY | 0.3623 | 1.5382 | 0.9674 | 2.9760 | 1.8321 <-E |
| 569 | DIQTVVKEY | 0.1160 | 0.4924 | 0.9665 | 2.5380 | 0.7643 <-E |
| 573 | VVKEYNERY | 0.1967 | 0.8353 | 0.9655 | 3.1330 | 1.1368 <-E |
| 599 | DIDTVVREY | 0.4725 | 2.0063 | 0.9535 | 2.3940 | 2.2690 <-E |
| 621 | SSPKPDPLY | 0.2237 | 0.9499 | 0.9712 | 3.0570 | 1.2485 <-E |
| 644 | DIVTKQSDY | 0.1120 | 0.4754 | 0.9624 | 2.8620 | 0.7629 <-E |
| 767 | QSKPNDDTY | 0.2887 | 1.2260 | 0.9706 | 2.9850 | 1.5208 <-E |
| 813 | TMSKISTKF | 0.1118 | 0.4747 | 0.9745 | 2.6850 | 0.7551 <-E |
| 899 | IIDTIKDIY | 0.5021 | 2.1320 | 0.7047 | 2.8700 | 2.3812 <-E |
| 931 | MSDNNKMGV | 0.3502 | 1.4867 | 0.9273 | 0.2740 | 1.6395 <-E |
| 963 | QNTDAMALY | 0.1404 | 0.5962 | 0.9208 | 2.8500 | 0.8769 <-E |
| 964 | NTDAMALYF | 0.7012 | 2.9773 | 0.2818 | 2.3250 | 3.1359 <-E |
| 972 | FLDVIDSEI | 0.1719 | 0.7298 | 0.5389 | 0.2310 | 0.8221 <-E |
| 974 | DVIDSEILY | 0.1294 | 0.5494 | 0.9420 | 2.8640 | 0.8339 <-E |
| 975 | VIDSEILYL | 0.1505 | 0.6388 | 0.8876 | 0.8300 | 0.8135 <-E |
| 982 | YLNTSNLVL | 0.1464 | 0.6215 | 0.9097 | 0.9080 | 0.8034 <-E |
| 984 | NTSNLVLEY | 0.7968 | 3.3830 | 0.9685 | 2.9840 | 3.6774 <-E |
| 1005 | SVDVDITAY | 0.6239 | 2.6490 | 0.9691 | 3.0720 | 2.9479 <-E |
| 1009 | DITAYTILY | 0.3332 | 1.4148 | 0.9740 | 2.7480 | 1.6983 <-E |
| 1018 | DTADNIKKY | 0.3513 | 1.4917 | 0.9417 | 2.5520 | 1.7606 <-E |
| 1060 | ISDMQLLKM | 0.2133 | 0.9057 | 0.9513 | 0.0470 | 1.0507 <-E |
| 1094 | YLLAGGCPY | 0.1462 | 0.6206 | 0.9726 | 2.8370 | 0.9084 <-E |
| 1111 | HTTCSILLR | 0.1404 | 0.5959 | 0.6701 | 1.3830 | 0.7656 <-E |
| 1192 | CSSRTRKIY | 0.2490 | 1.0571 | 0.1648 | 2.9740 | 1.2305 <-E |
| 1238 | IIELPVGDY | 0.1955 | 0.8299 | 0.9433 | 2.9790 | 1.1203 <-E |
| 1252 | YSATKPSRI | 0.1634 | 0.6938 | 0.2680 | 0.6200 | 0.7650 <-E |
| 1260 | IAVFCTHNY | 0.1058 | 0.4492 | 0.9573 | 3.2100 | 0.7533 <-E |
| 1273 | KSDIIVLMF | 0.5623 | 2.3873 | 0.9337 | 2.4010 | 2.6474 <-E |
| 1336 | DSVETDIHY | 0.2547 | 1.0813 | 0.8074 | 2.5600 | 1.3304 <-E |
| 1380 | MVDNLGNGY | 0.6877 | 2.9200 | 0.8658 | 2.8290 | 3.1914 <-E |
| 1437 | YLQSTSQDY | 0.3379 | 1.4345 | 0.9011 | 2.9390 | 1.7167 <-E |
| 1482 | SIPRNISTY | 0.1287 | 0.5464 | 0.9758 | 2.9860 | 0.8420 <-E |
| 1525 | RACFHHWNY | 0.3270 | 1.3883 | 0.8099 | 3.0850 | 1.6641 <-E |
| 1532 | NYYTLSLDY | 0.1121 | 0.4761 | 0.9718 | 3.3570 | 0.7897 <-E |
| 1533 | YYTLSLDYY | 0.1099 | 0.4666 | 0.9469 | 3.2740 | 0.7724 <-E |
| 1536 | LSLDYYCSY | 0.4286 | 1.8198 | 0.8891 | 2.9970 | 2.1030 <-E |
| 1558 | CKSYIHIEY | 0.1989 | 0.8446 | 0.9262 | 2.7840 | 1.1227 <-E |
| 1585 | FIHDNSNEY | 0.3652 | 1.5507 | 0.9708 | 2.8210 | 1.8374 <-E |
| 1598 | ISNKLNDLY | 0.6218 | 2.6403 | 0.4412 | 2.7270 | 2.8428 <-E |
| 1601 | KLNDLYNEY | 0.3068 | 1.3026 | 0.9555 | 3.0980 | 1.6008 <-E |
| 1697 | IIDVKNNLV | 0.1916 | 0.8135 | 0.5823 | 0.2830 | 0.9150 <-E |
| 1714 | YLCDTLDDY | 0.3048 | 1.2941 | 0.6966 | 3.0740 | 1.5523 <-E |
| 1725 | TSVEYNKSY | 0.2700 | 1.1465 | 0.9560 | 2.9700 | 1.4384 <-E |
| 1735 | LVNDTYISY | 0.3238 | 1.3748 | 0.9775 | 3.0200 | 1.6724 <-E |
| 1808 | YLNSSPDIY | 0.4086 | 1.7347 | 0.4269 | 2.9710 | 1.9472 <-E |
| 1812 | SPDIYHIIY | 0.2862 | 1.2153 | 0.9699 | 2.5510 | 1.4883 <-E |
| 1872 | SMEIIDNRY | 0.5615 | 2.3839 | 0.9756 | 3.0020 | 2.6803 <-E |
| 2023-MPXV-B21R | | | | | | |
| 12 | LTVTCSWCY | 0.6345 | 2.6940 | 0.9382 | 2.9330 | 2.9814 <-E |
| 44 | SVASLPYKY | 0.2833 | 1.2029 | 0.7983 | 2.9940 | 1.4723 <-E |
| 70 | WTDIAEGVR | 0.1693 | 0.7187 | 0.0432 | 1.4010 | 0.7952 <-E |
| 83 | KICDINGTY | 0.3071 | 1.3038 | 0.8688 | 3.1780 | 1.5930 <-E |
| 110 | PTVTPITTY | 0.2319 | 0.9844 | 0.9723 | 2.5490 | 1.2577 <-E |
| 115 | ITTYEPSIY | 0.4981 | 2.1148 | 0.6249 | 2.7450 | 2.3458 <-E |
| 117 | TYEPSIYNY | 0.1394 | 0.5919 | 0.9741 | 3.0340 | 0.8897 <-E |
| 121 | SIYNYTIDY | 0.2145 | 0.9106 | 0.9727 | 3.1480 | 1.2139 <-E |
| 182 | ETELTNYLI | 0.2056 | 0.8731 | 0.6316 | 0.3060 | 0.9831 <-E |
| 206 | ETSNNNTLH | 0.2680 | 1.1380 | 0.1142 | -0.9040 | 1.1100 <-E |
| 242 | DTSEEILLM | 0.2115 | 0.8980 | 0.8967 | 0.1040 | 1.0377 <-E |
| 243 | TSEEILLML | 0.2014 | 0.8550 | 0.9652 | 0.6910 | 1.0343 <-E |
| 256 | SSDTFISST | 0.2687 | 1.1406 | 0.3474 | -0.7340 | 1.1561 <-E |
| 266 | ITECLKTLI | 0.1650 | 0.7005 | 0.2384 | 0.5330 | 0.7629 <-E |
| 322 | KDDENNTVY | 0.1514 | 0.6429 | 0.9539 | 2.5950 | 0.9157 <-E |
| 358 | LTNIIHNSV | 0.1606 | 0.6820 | 0.9502 | 0.2130 | 0.8352 <-E |
| 385 | TSKELDCLY | 0.4246 | 1.8030 | 0.8142 | 2.9960 | 2.0749 <-E |
| 388 | ELDCLYESY | 0.5389 | 2.2882 | 0.9343 | 2.6650 | 2.5616 <-E |
| 413 | RSDDKKEYM | 0.1490 | 0.6327 | 0.6499 | 0.5450 | 0.7575 <-E |
| 420 | YMDMKLFDH | 0.1853 | 0.7867 | 0.1055 | -0.7770 | 0.7637 <-E |
| 484 | ITATEADLY | 0.7400 | 3.1419 | 0.5413 | 2.7980 | 3.3630 <-E |
| 510 | FTEALVSTI | 0.2466 | 1.0469 | 0.3170 | 0.1290 | 1.1009 <-E |
| 522 | LSNVREVTY | 0.3623 | 1.5382 | 0.9674 | 2.9760 | 1.8321 <-E |
| 569 | DIQTVVKEY | 0.1160 | 0.4924 | 0.9665 | 2.5380 | 0.7643 <-E |
| 573 | VVKEYNERY | 0.1967 | 0.8353 | 0.9655 | 3.1330 | 1.1368 <-E |
| 599 | DIDTVVREY | 0.4725 | 2.0063 | 0.9535 | 2.3940 | 2.2690 <-E |
| 621 | SSPKPDPLY | 0.2237 | 0.9499 | 0.9712 | 3.0570 | 1.2485 <-E |
| 644 | DIVTKQSDY | 0.1120 | 0.4754 | 0.9624 | 2.8620 | 0.7629 <-E |
| 767 | QSKPNDDTY | 0.2887 | 1.2260 | 0.9706 | 2.9850 | 1.5208 <-E |
| 813 | TMSKISTKF | 0.1118 | 0.4747 | 0.9745 | 2.6850 | 0.7551 <-E |
| 899 | IIDTIKDIY | 0.5021 | 2.1320 | 0.7047 | 2.8700 | 2.3812 <-E |
| 931 | MSDNNKMGV | 0.3502 | 1.4867 | 0.9273 | 0.2740 | 1.6395 <-E |
| 963 | QNTDAMALY | 0.1404 | 0.5962 | 0.9208 | 2.8500 | 0.8769 <-E |
| 964 | NTDAMALYF | 0.7012 | 2.9773 | 0.2818 | 2.3250 | 3.1359 <-E |
| 972 | FLDVIDSEI | 0.1719 | 0.7298 | 0.5389 | 0.2310 | 0.8221 <-E |
| 974 | DVIDSEILY | 0.1294 | 0.5494 | 0.9420 | 2.8640 | 0.8339 <-E |
| 975 | VIDSEILYL | 0.1505 | 0.6388 | 0.8876 | 0.8300 | 0.8135 <-E |
| 982 | YLNTSNLVL | 0.1464 | 0.6215 | 0.9097 | 0.9080 | 0.8034 <-E |
| 984 | NTSNLVLEY | 0.7968 | 3.3830 | 0.9685 | 2.9840 | 3.6774 <-E |
| 1005 | SVDVDITAY | 0.6239 | 2.6490 | 0.9691 | 3.0720 | 2.9479 <-E |
| 1009 | DITAYTILY | 0.3332 | 1.4148 | 0.9740 | 2.7480 | 1.6983 <-E |
| 1018 | DTADNIKKY | 0.3513 | 1.4917 | 0.9417 | 2.5520 | 1.7606 <-E |
| 1060 | ISDMQLLKM | 0.2133 | 0.9057 | 0.9513 | 0.0470 | 1.0507 <-E |
| 1094 | YLLAGGCPY | 0.1462 | 0.6206 | 0.9726 | 2.8370 | 0.9084 <-E |
| 1111 | HTTCSILLR | 0.1404 | 0.5959 | 0.6701 | 1.3830 | 0.7656 <-E |
| 1192 | CSSRTRKIY | 0.2490 | 1.0571 | 0.1648 | 2.9740 | 1.2305 <-E |
| 1238 | IIELPVGDY | 0.1955 | 0.8299 | 0.9433 | 2.9790 | 1.1203 <-E |
| 1252 | YSATKPSRI | 0.1634 | 0.6938 | 0.2680 | 0.6200 | 0.7650 <-E |
| 1260 | IAVFCTHNY | 0.1058 | 0.4492 | 0.9573 | 3.2100 | 0.7533 <-E |
| 1273 | KSDIIVLMF | 0.5623 | 2.3873 | 0.9337 | 2.4010 | 2.6474 <-E |
| 1336 | DSVETDIHY | 0.2547 | 1.0813 | 0.8074 | 2.5600 | 1.3304 <-E |
| 1380 | MVDNLGNGY | 0.6877 | 2.9200 | 0.8658 | 2.8290 | 3.1914 <-E |
| 1437 | YLQSTSQDY | 0.3379 | 1.4345 | 0.9011 | 2.9390 | 1.7167 <-E |
| 1482 | SIPRNISTY | 0.1287 | 0.5464 | 0.9758 | 2.9860 | 0.8420 <-E |
| 1525 | RACFHHWNY | 0.3270 | 1.3883 | 0.8099 | 3.0850 | 1.6641 <-E |
| 1532 | NYYTLSLDY | 0.1121 | 0.4761 | 0.9718 | 3.3570 | 0.7897 <-E |
| 1533 | YYTLSLDYY | 0.1099 | 0.4666 | 0.9469 | 3.2740 | 0.7724 <-E |
| 1536 | LSLDYYCSY | 0.4286 | 1.8198 | 0.8891 | 2.9970 | 2.1030 <-E |
| 1558 | CKSYIHIEY | 0.1989 | 0.8446 | 0.9262 | 2.7840 | 1.1227 <-E |
| 1585 | FIHDNSNEY | 0.3652 | 1.5507 | 0.9708 | 2.8210 | 1.8374 <-E |
| 1598 | ISNKLNDLY | 0.6218 | 2.6403 | 0.4412 | 2.7270 | 2.8428 <-E |
| 1601 | KLNDLYNEY | 0.3068 | 1.3026 | 0.9555 | 3.0980 | 1.6008 <-E |
| 1697 | IIDVKNNLV | 0.1916 | 0.8135 | 0.5823 | 0.2830 | 0.9150 <-E |
| 1714 | YLCDTLDDY | 0.3048 | 1.2941 | 0.6966 | 3.0740 | 1.5523 <-E |
| 1725 | TSVEYNKSY | 0.2700 | 1.1465 | 0.9560 | 2.9700 | 1.4384 <-E |
| 1735 | LVNDTYISY | 0.3238 | 1.3748 | 0.9775 | 3.0200 | 1.6724 <-E |
| 1808 | YLNSSPDIY | 0.4086 | 1.7347 | 0.4269 | 2.9710 | 1.9472 <-E |
| 1812 | SPDIYHIIY | 0.2862 | 1.2153 | 0.9699 | 2.5510 | 1.4883 <-E |
| 1872 | SMEIIDNRY | 0.5615 | 2.3839 | 0.9756 | 3.0020 | 2.6803 <-E |
| 2024-MPXV-B21R | | | | | | |
| 12 | LTVTCSWCY | 0.6345 | 2.6940 | 0.9382 | 2.9330 | 2.9814 <-E |
| 44 | SVASLPYKY | 0.2833 | 1.2029 | 0.7983 | 2.9940 | 1.4723 <-E |
| 70 | WTDIAEGVR | 0.1693 | 0.7187 | 0.0432 | 1.4010 | 0.7952 <-E |
| 83 | KICDINGTY | 0.3071 | 1.3038 | 0.8688 | 3.1780 | 1.5930 <-E |
| 110 | PTVTPITTY | 0.2319 | 0.9844 | 0.9723 | 2.5490 | 1.2577 <-E |
| 115 | ITTYEPSIY | 0.4981 | 2.1148 | 0.6249 | 2.7450 | 2.3458 <-E |
| 117 | TYEPSIYNY | 0.1394 | 0.5919 | 0.9741 | 3.0340 | 0.8897 <-E |
| 121 | SIYNYTIDY | 0.2145 | 0.9106 | 0.9727 | 3.1480 | 1.2139 <-E |
| 182 | ETELTNYLI | 0.2056 | 0.8731 | 0.6316 | 0.3060 | 0.9831 <-E |
| 206 | ETSNNNTLH | 0.2680 | 1.1380 | 0.1142 | -0.9040 | 1.1100 <-E |
| 242 | DTSEEILLM | 0.2115 | 0.8980 | 0.8967 | 0.1040 | 1.0377 <-E |
| 243 | TSEEILLML | 0.2014 | 0.8550 | 0.9652 | 0.6910 | 1.0343 <-E |
| 256 | SSDTFISST | 0.2687 | 1.1406 | 0.3474 | -0.7340 | 1.1561 <-E |
| 266 | ITECLKTLI | 0.1650 | 0.7005 | 0.2384 | 0.5330 | 0.7629 <-E |
| 322 | KDDENNTVY | 0.1514 | 0.6429 | 0.9539 | 2.5950 | 0.9157 <-E |
| 358 | LTNIIHNSV | 0.1606 | 0.6820 | 0.9502 | 0.2130 | 0.8352 <-E |
| 385 | TSKELDCLY | 0.4246 | 1.8030 | 0.8142 | 2.9960 | 2.0749 <-E |
| 388 | ELDCLYESY | 0.5389 | 2.2882 | 0.9343 | 2.6650 | 2.5616 <-E |
| 413 | RSDDKKEYM | 0.1490 | 0.6327 | 0.6499 | 0.5450 | 0.7575 <-E |
| 420 | YMDMKLFDH | 0.1853 | 0.7867 | 0.1055 | -0.7770 | 0.7637 <-E |
| 484 | ITATEADLY | 0.7400 | 3.1419 | 0.5413 | 2.7980 | 3.3630 <-E |
| 510 | FTEALVSTI | 0.2466 | 1.0469 | 0.3170 | 0.1290 | 1.1009 <-E |
| 522 | LSNVREVTY | 0.3623 | 1.5382 | 0.9674 | 2.9760 | 1.8321 <-E |
| 569 | DIQTVVKEY | 0.1160 | 0.4924 | 0.9665 | 2.5380 | 0.7643 <-E |
| 573 | VVKEYNERY | 0.1967 | 0.8353 | 0.9655 | 3.1330 | 1.1368 <-E |
| 599 | DIDTVVREY | 0.4725 | 2.0063 | 0.9535 | 2.3940 | 2.2690 <-E |
| 621 | SSPKPDPLY | 0.2237 | 0.9499 | 0.9712 | 3.0570 | 1.2485 <-E |
| 644 | DIVTKQSDY | 0.1120 | 0.4754 | 0.9624 | 2.8620 | 0.7629 <-E |
| 767 | QSKPNDDTY | 0.2887 | 1.2260 | 0.9706 | 2.9850 | 1.5208 <-E |
| 813 | TMSKISTKF | 0.1118 | 0.4747 | 0.9745 | 2.6850 | 0.7551 <-E |
| 899 | IIDTIKDIY | 0.5021 | 2.1320 | 0.7047 | 2.8700 | 2.3812 <-E |
| 931 | MSDNNKMGV | 0.3502 | 1.4867 | 0.9273 | 0.2740 | 1.6395 <-E |
| 963 | QNTDAMALY | 0.1404 | 0.5962 | 0.9208 | 2.8500 | 0.8769 <-E |
| 964 | NTDAMALYF | 0.7012 | 2.9773 | 0.2818 | 2.3250 | 3.1359 <-E |
| 972 | FLDVIDSEI | 0.1719 | 0.7298 | 0.5389 | 0.2310 | 0.8221 <-E |
| 974 | DVIDSEILY | 0.1294 | 0.5494 | 0.9420 | 2.8640 | 0.8339 <-E |
| 975 | VIDSEILYL | 0.1505 | 0.6388 | 0.8876 | 0.8300 | 0.8135 <-E |
| 982 | YLNTSNLVL | 0.1464 | 0.6215 | 0.9097 | 0.9080 | 0.8034 <-E |
| 984 | NTSNLVLEY | 0.7968 | 3.3830 | 0.9685 | 2.9840 | 3.6774 <-E |
| 1005 | SVDVDITAY | 0.6239 | 2.6490 | 0.9691 | 3.0720 | 2.9479 <-E |
| 1009 | DITAYTILY | 0.3332 | 1.4148 | 0.9740 | 2.7480 | 1.6983 <-E |
| 1018 | DTADNIKKY | 0.3513 | 1.4917 | 0.9417 | 2.5520 | 1.7606 <-E |
| 1060 | ISDMQLLKM | 0.2133 | 0.9057 | 0.9513 | 0.0470 | 1.0507 <-E |
| 1094 | YLLAGGCPY | 0.1462 | 0.6206 | 0.9726 | 2.8370 | 0.9084 <-E |
| 1111 | HTTCSILLR | 0.1404 | 0.5959 | 0.6701 | 1.3830 | 0.7656 <-E |
| 1192 | CSSRTRKIY | 0.2490 | 1.0571 | 0.1648 | 2.9740 | 1.2305 <-E |
| 1238 | IIELPVGDY | 0.1955 | 0.8299 | 0.9433 | 2.9790 | 1.1203 <-E |
| 1252 | YSATKPSRI | 0.1634 | 0.6938 | 0.2680 | 0.6200 | 0.7650 <-E |
| 1260 | IAVFCTHNY | 0.1058 | 0.4492 | 0.9573 | 3.2100 | 0.7533 <-E |
| 1273 | KSDIIVLMF | 0.5623 | 2.3873 | 0.9337 | 2.4010 | 2.6474 <-E |
| 1336 | DSVETDIHY | 0.2547 | 1.0813 | 0.8074 | 2.5600 | 1.3304 <-E |
| 1380 | MVDNLGNGY | 0.6877 | 2.9200 | 0.8658 | 2.8290 | 3.1914 <-E |
| 1437 | YLQSTSQDY | 0.3379 | 1.4345 | 0.9011 | 2.9390 | 1.7167 <-E |
| 1482 | SIPRNISTY | 0.1287 | 0.5464 | 0.9758 | 2.9860 | 0.8420 <-E |
| 1525 | RACFHHWNY | 0.3270 | 1.3883 | 0.8099 | 3.0850 | 1.6641 <-E |
| 1532 | NYYTLSLDY | 0.1121 | 0.4761 | 0.9718 | 3.3570 | 0.7897 <-E |
| 1533 | YYTLSLDYY | 0.1099 | 0.4666 | 0.9469 | 3.2740 | 0.7724 <-E |
| 1536 | LSLDYYCSY | 0.4286 | 1.8198 | 0.8891 | 2.9970 | 2.1030 <-E |
| 1558 | CKSYIHIEY | 0.1989 | 0.8446 | 0.9262 | 2.7840 | 1.1227 <-E |
| 1585 | FIHDNSNEY | 0.3652 | 1.5507 | 0.9708 | 2.8210 | 1.8374 <-E |
| 1598 | ISNKLNDLY | 0.6218 | 2.6403 | 0.4412 | 2.7270 | 2.8428 <-E |
| 1601 | KLNDLYNEY | 0.3068 | 1.3026 | 0.9555 | 3.0980 | 1.6008 <-E |
| 1697 | IIDVKNNLV | 0.1916 | 0.8135 | 0.5823 | 0.2830 | 0.9150 <-E |
| 1714 | YLCDTLDDY | 0.3048 | 1.2941 | 0.6966 | 3.0740 | 1.5523 <-E |
| 1725 | TSVEYNKSY | 0.2700 | 1.1465 | 0.9560 | 2.9700 | 1.4384 <-E |
| 1735 | LVNDTYISY | 0.3238 | 1.3748 | 0.9775 | 3.0200 | 1.6724 <-E |
| 1808 | YLNSSPDIY | 0.4086 | 1.7347 | 0.4269 | 2.9710 | 1.9472 <-E |
| 1812 | SPDIYHIIY | 0.2862 | 1.2153 | 0.9699 | 2.5510 | 1.4883 <-E |
| 1872 | SMEIIDNRY | 0.5615 | 2.3839 | 0.9756 | 3.0020 | 2.6803 <-E |
| 2025-MPXV-B21R | | | | | | |
| 12 | LTVTCSWCY | 0.6345 | 2.6940 | 0.9382 | 2.9330 | 2.9814 <-E |
| 44 | SVASLPYKY | 0.2833 | 1.2029 | 0.7983 | 2.9940 | 1.4723 <-E |
| 70 | WTDIAEGVR | 0.1693 | 0.7187 | 0.0432 | 1.4010 | 0.7952 <-E |
| 83 | KICDINGTY | 0.3071 | 1.3038 | 0.8688 | 3.1780 | 1.5930 <-E |
| 110 | PTVTPITTY | 0.2319 | 0.9844 | 0.9723 | 2.5490 | 1.2577 <-E |
| 115 | ITTYEPSIY | 0.4981 | 2.1148 | 0.6249 | 2.7450 | 2.3458 <-E |
| 117 | TYEPSIYNY | 0.1394 | 0.5919 | 0.9741 | 3.0340 | 0.8897 <-E |
| 121 | SIYNYTIDY | 0.2145 | 0.9106 | 0.9727 | 3.1480 | 1.2139 <-E |
| 182 | ETELTNYLI | 0.2056 | 0.8731 | 0.6316 | 0.3060 | 0.9831 <-E |
| 206 | ETSDNNTLH | 0.2687 | 1.1410 | 0.1596 | -0.9040 | 1.1197 <-E |
| 207 | TSDNNTLHG | 0.2184 | 0.9274 | 0.1116 | -1.7190 | 0.8582 <-E |
| 242 | DTSEEILLM | 0.2115 | 0.8980 | 0.8967 | 0.1040 | 1.0377 <-E |
| 243 | TSEEILLML | 0.2014 | 0.8550 | 0.9652 | 0.6910 | 1.0343 <-E |
| 256 | SSDTFISST | 0.2687 | 1.1406 | 0.3474 | -0.7340 | 1.1561 <-E |
| 266 | ITECLKTLI | 0.1650 | 0.7005 | 0.2384 | 0.5330 | 0.7629 <-E |
| 322 | KDDENNTVY | 0.1514 | 0.6429 | 0.9539 | 2.5950 | 0.9157 <-E |
| 358 | LTNIIHNSV | 0.1606 | 0.6820 | 0.9502 | 0.2130 | 0.8352 <-E |
| 385 | TSKELDCLY | 0.4246 | 1.8030 | 0.8142 | 2.9960 | 2.0749 <-E |
| 388 | ELDCLYESY | 0.5389 | 2.2882 | 0.9343 | 2.6650 | 2.5616 <-E |
| 413 | RSDDKKEYM | 0.1490 | 0.6327 | 0.6499 | 0.5450 | 0.7575 <-E |
| 420 | YMDMKLFDH | 0.1853 | 0.7867 | 0.1055 | -0.7770 | 0.7637 <-E |
| 484 | ITATEADLY | 0.7400 | 3.1419 | 0.5413 | 2.7980 | 3.3630 <-E |
| 510 | FTEALVSTI | 0.2466 | 1.0469 | 0.3170 | 0.1290 | 1.1009 <-E |
| 522 | LSNVREVTY | 0.3623 | 1.5382 | 0.9674 | 2.9760 | 1.8321 <-E |
| 569 | DIQTVVKEY | 0.1160 | 0.4924 | 0.9665 | 2.5380 | 0.7643 <-E |
| 573 | VVKEYNERY | 0.1967 | 0.8353 | 0.9655 | 3.1330 | 1.1368 <-E |
| 599 | DIDTVVREY | 0.4725 | 2.0063 | 0.9535 | 2.3940 | 2.2690 <-E |
| 621 | SSPKPDPLY | 0.2237 | 0.9499 | 0.9712 | 3.0570 | 1.2485 <-E |
| 644 | DIVTKQSDY | 0.1120 | 0.4754 | 0.9624 | 2.8620 | 0.7629 <-E |
| 767 | QSKPNDDTY | 0.2887 | 1.2260 | 0.9706 | 2.9850 | 1.5208 <-E |
| 813 | TMSKISTKF | 0.1118 | 0.4747 | 0.9745 | 2.6850 | 0.7551 <-E |
| 899 | IIDTIKDIY | 0.5021 | 2.1320 | 0.7047 | 2.8700 | 2.3812 <-E |
| 931 | MSDNNKMGV | 0.3502 | 1.4867 | 0.9273 | 0.2740 | 1.6395 <-E |
| 963 | QNTDAMALY | 0.1404 | 0.5962 | 0.9208 | 2.8500 | 0.8769 <-E |
| 964 | NTDAMALYF | 0.7012 | 2.9773 | 0.2818 | 2.3250 | 3.1359 <-E |
| 972 | FLDVIDSEI | 0.1719 | 0.7298 | 0.5389 | 0.2310 | 0.8221 <-E |
| 974 | DVIDSEILY | 0.1294 | 0.5494 | 0.9420 | 2.8640 | 0.8339 <-E |
| 975 | VIDSEILYL | 0.1505 | 0.6388 | 0.8876 | 0.8300 | 0.8135 <-E |
| 982 | YLNTSNLVL | 0.1464 | 0.6215 | 0.9097 | 0.9080 | 0.8034 <-E |
| 984 | NTSNLVLEY | 0.7968 | 3.3830 | 0.9685 | 2.9840 | 3.6774 <-E |
| 1005 | SVDVDITAY | 0.6239 | 2.6490 | 0.9691 | 3.0720 | 2.9479 <-E |
| 1009 | DITAYTILY | 0.3332 | 1.4148 | 0.9740 | 2.7480 | 1.6983 <-E |
| 1018 | DTADNIKKY | 0.3513 | 1.4917 | 0.9417 | 2.5520 | 1.7606 <-E |
| 1060 | ISDMQLLKM | 0.2133 | 0.9057 | 0.9513 | 0.0470 | 1.0507 <-E |
| 1094 | YLLAGGCPY | 0.1462 | 0.6206 | 0.9726 | 2.8370 | 0.9084 <-E |
| 1111 | HTTCSILLR | 0.1404 | 0.5959 | 0.6701 | 1.3830 | 0.7656 <-E |
| 1192 | CSSRTRKIY | 0.2490 | 1.0571 | 0.1648 | 2.9740 | 1.2305 <-E |
| 1238 | IIELPVGDY | 0.1955 | 0.8299 | 0.9433 | 2.9790 | 1.1203 <-E |
| 1252 | YSATKPSRI | 0.1634 | 0.6938 | 0.2680 | 0.6200 | 0.7650 <-E |
| 1260 | IAVFCTHNY | 0.1058 | 0.4492 | 0.9573 | 3.2100 | 0.7533 <-E |
| 1273 | KSDIIVLMF | 0.5623 | 2.3873 | 0.9337 | 2.4010 | 2.6474 <-E |
| 1336 | DSVETDIHY | 0.2547 | 1.0813 | 0.8074 | 2.5600 | 1.3304 <-E |
| 1380 | MVDNLGNGY | 0.6877 | 2.9200 | 0.8658 | 2.8290 | 3.1914 <-E |
| 1437 | YLQSTSQDY | 0.3379 | 1.4345 | 0.9011 | 2.9390 | 1.7167 <-E |
| 1482 | SIPRNISTY | 0.1287 | 0.5464 | 0.9758 | 2.9860 | 0.8420 <-E |
| 1525 | RACFHHWNY | 0.3270 | 1.3883 | 0.8099 | 3.0850 | 1.6641 <-E |
| 1532 | NYYTLSLDY | 0.1121 | 0.4761 | 0.9718 | 3.3570 | 0.7897 <-E |
| 1533 | YYTLSLDYY | 0.1099 | 0.4666 | 0.9469 | 3.2740 | 0.7724 <-E |
| 1536 | LSLDYYCSY | 0.4286 | 1.8198 | 0.8891 | 2.9970 | 2.1030 <-E |
| 1558 | CKSYIHIEY | 0.1989 | 0.8446 | 0.9262 | 2.7840 | 1.1227 <-E |
| 1585 | FIHDNSNEY | 0.3652 | 1.5507 | 0.9708 | 2.8210 | 1.8374 <-E |
| 1598 | ISNKLNDLY | 0.6218 | 2.6403 | 0.4412 | 2.7270 | 2.8428 <-E |
| 1601 | KLNDLYNEY | 0.3068 | 1.3026 | 0.9555 | 3.0980 | 1.6008 <-E |
| 1697 | IIDVKNNLV | 0.1916 | 0.8135 | 0.5823 | 0.2830 | 0.9150 <-E |
| 1714 | YLCDTLDDY | 0.3048 | 1.2941 | 0.6966 | 3.0740 | 1.5523 <-E |
| 1725 | TSVEYNKSY | 0.2700 | 1.1465 | 0.9560 | 2.9700 | 1.4384 <-E |
| 1735 | LVNDTYISY | 0.3238 | 1.3748 | 0.9775 | 3.0200 | 1.6724 <-E |
| 1808 | YLNSSPDIY | 0.4086 | 1.7347 | 0.4269 | 2.9710 | 1.9472 <-E |
| 1812 | SPDIYHIIY | 0.2862 | 1.2153 | 0.9699 | 2.5510 | 1.4883 <-E |
| 1872 | SMEIINNRY | 0.6439 | 2.7338 | 0.9772 | 3.0020 | 3.0304 <-E |

| No. | Chain | Start | End | Peptide | Residues | Score |
| --- | --- | --- | --- | --- | --- | --- |
| Zaire-96-I-16-B21R _(1-300)_ | | | | | | |
| 1 | A | 117 | 164 | TYEPSIYNYTIDYSTVITTEELQVTPTYAPVTTPLPTSAVPYDQRSNN | 48 | 0.804 |
| 2 | A | 14 | 76 | VTCSWCYETCMRKTALYHDIQLEHVEDNKDSVASLPYKYLQVVKQRERSRLLATFNWTDIAEG | 63 | 0.778 |
| 3 | A | 196 | 211 | TVDNNTMVDDETSDNN | 16 | 0.65 |
| 4 | A | 86 | 91 | DINGTY | 6 | 0.569 |
| 5 | A | 279 | 283 | INDVL | 5 | 0.546 |
| 6 | A | 104 | 109 | DSTEEL | 6 | 0.533 |
| 2003-MPXV-B21R _(1-300)_ | | | | | | |
| 1 | A | 117 | 162 | TYEPSIYNYTIDYSTVITTEELQVTPTYAPVTTPLPTSAVPYDQRS | 46 | 0.776 |
| 2 | A | 28 | 76 | ALYHDIQLEHVEDNKDSVASLPYKYLQVVKQRERSRLLATFNWTDIAEG | 49 | 0.755 |
| 3 | A | 1 | 12 | MNFQKLSLAIYL | 12 | 0.731 |
| 4 | A | 275 | 285 | NNVSINDVLIT | 11 | 0.631 |
| 5 | A | 102 | 109 | IIDSTEEL | 8 | 0.599 |
| 6 | A | 199 | 210 | NNTMVDDETSDN | 12 | 0.591 |
| 7 | A | 86 | 91 | DINGTY | 6 | 0.533 |
| 2017-MPXV-B21R _(1-300)_ | | | | | | |
| 1 | A | 106 | 154 | TEELPTVTPITTYEPSIYNYTIDYSTVITTEELQVTPTYAPVTTPLPTS | 49 | 0.821 |
| 2 | A | 27 | 91 | TALYHDIQLEHVEDNKDSVASLPYKYLQVVKQRERSRLLATFNWTDIAEGVRNEFIKICDINGTY | 65 | 0.712 |
| 3 | A | 194 | 211 | NDTVDNNTMVDDETSDNN | 18 | 0.697 |
| 2022-MPXV-B21R _(1-300)_ | | | | | | |
| 1 | A | 104 | 154 | DSTEELPTVTPITTYEPSIYNYTIDYSTVITTEELQVTPTYAPVTTPLPTS | 51 | 0.808 |
| 2 | A | 1 | 13 | MNFQKLSLAIYLT | 13 | 0.733 |
| 3 | A | 28 | 91 | ALYHDIQLEHVEDNKDSVASLPYKYLQVVKQRERSRLLATFNWTDIAEGVRNEFIKICDINGTY | 64 | 0.708 |
| 4 | A | 193 | 211 | KNDTVDNNTMVDDETSNNN | 19 | 0.644 |
| 5 | A | 279 | 283 | INDVL | 5 | 0.502 |
| 2023-MPXV-B21R _(1-300)_ | | | | | | |
| 1 | A | 110 | 152 | PTVTPITTYEPSIYNYTIDYSTVITTEELQVTPTYAPVTTPLP | 43 | 0.816 |
| 2 | A | 1 | 16 | MNFQKLSLAIYLTVTC | 16 | 0.807 |
| 3 | A | 194 | 211 | NDTVDNNTMVDDETSNNN | 18 | 0.71 |
| 4 | A | 26 | 93 | KTALYHDIQLEHVEDNKDSVASLPYKYLQVVKQRERSRLLATFNWTDIAEGVRNEFIKICDINGTYLY | 68 | 0.693 |
| 5 | A | 257 | 265 | SDTFISSTN | 9 | 0.533 |
| 2024-MPXV-B21R _(1-300)_ | | | | | | |
| 1 | A | 1 | 13 | MNFQKLSLAIYLT | 13 | 0.773 |
| 2 | A | 104 | 144 | DSTEELPTVTPITTYEPSIYNYTIDYSTVITTEELQVTPTY | 41 | 0.741 |
| 3 | A | 29 | 91 | LYHDIQLEHVEDNKDSVASLPYKYLQVVKQRERSRLLATFNWTDIAEGVRNEFIKICDINGTY | 63 | 0.719 |
| 4 | A | 194 | 211 | NDTVDNNTMVDDETSNNN | 18 | 0.686 |
| 5 | A | 154 | 166 | SAVPYDQRSNNNV | 13 | 0.606 |
| 2025-MPXV-B21R _(1-300)_ | | | | | | |
| 1 | A | 33 | 80 | IQLEHVEDNKDSVASLPYKYLQVVKQRERSRLLATFNWTDIAEGVRNE | 48 | 0.788 |
| 2 | A | 113 | 164 | TPITTYEPSIYNYTIDYSTVITTEELQVTPTYAPVTTPLPTSAVPYDQRSNN | 52 | 0.748 |
| 3 | A | 1 | 10 | MNFQKLSLAI | 10 | 0.722 |
| 4 | A | 194 | 211 | NDTVDNNTMVDDETSDNN | 18 | 0.691 |
| 5 | A | 104 | 110 | DSTEELP | 7 | 0.639 |
| 6 | A | 275 | 285 | NNVSINDVLIT | 11 | 0.571 |
| 7 | A | 83 | 91 | KICDINGTY | 9 | 0.527 |

Table S7. B cell epitope based on struc
